# Multicellular group formation as a resource competition strategy for marine bacteria

**DOI:** 10.1101/2025.08.01.668187

**Authors:** Thomas C. Day, Julia A. Schwartzman

## Abstract

Multicellularity can enhance bacterial resilience and robustness in nature. Traits like access to otherwise inaccessible substrates, increased stress tolerance and predation resistance provide ecological advantages that may scaffold the evolution of multicellular complexity. While multicellularity arises in many ecological contexts, we lack general principles to predict which environments favor these traits. Here, we ask how the physical size of multicellular groups affects their ability to compete for resources, focusing on marine bacteria that degrade organic matter. These bacteria compete for resources ranging from nanoscale dissolved organic matter to millimeter-scale particles. Using a theory of size-dependent fluid encounters, we identify a regime where resource encounters depend on group size. Experiments with the group-forming marine bacteria *Vibrio splendidus* 12B01 support the prediction that larger groups encounter resources more frequently. However, analysis of per-capita resource allocation reveals a trade-off: group size is inversely related to both the mean and variance of per-capita encounters. Cells in groups obtain fewer resources per cell than single cells, but experience more consistent encounters, sheltering them from large fluctuations in resource acquisition. Stochastic simulations support this tradeoff and predict a broad range of conditions in which consistency enables multicellular groups to out-compete or coexist with single cells. Our findings suggest that group formation is a potentially widespread but overlooked strategy for resource competition in marine heterotrophic bacteria, shaped by biophysical constraints on resource encounters.

**Significance Statement:** Multicellular groups are thought to arise in ecological settings where group-level traits confer a fitness advantage. However, linking these evolutionary drivers to specific ecological mechanisms remains a major challenge in understanding the origins of multicellular life. In this study, we draw inspiration from the microscale ecology of marine bacteria that play a key role in the ocean’s carbon cycle. These bacteria often rely on high local population densities to secrete enough extracellular enzymes to degrade organic matter and access nutrients. By combining experimental measurements with a mathematical theory of encounters and stochastic simulations of competition dynamics, we show that multicellular bacterial groups often out-compete single cells. Our results suggest that the advantage of multicellular groups stems from a tradeoff between the consistency and quantity of resource encounters, arising from purely physical processes. The physics of encounters offers a generic framework that can be leveraged to understand ecological scaffolds of multicellularity.

## Introduction

Multicellularity hides in plain sight in microbial ecosystems. The formation of multicellular groups - such as biofilms, bundles, and aggregates - can help microbes overcome ecological barriers that face single cells [1]. Beyond conspicuous structures like fruiting bodies and filaments, multicellular organization is a recurring motif in natural microbial communities [2–5]. Group-level traits like the metabolism of otherwise inaccessible substrates, increased resilience to abiotic stress, and resistance to predation are all potential ecological drivers for the evolution of multicellular groups [6]. By placing these drivers into an environmental context [7–9], we can better understand how the ecological properties of groups scaffold the evolution of multicellularity [10].

Marine environments concentrate many of the key ecological drivers of multicellularity. At the microscale, resources often exist in particulate form or are adsorbed onto particles, creating access challenges for individual cells [11]. In parallel, the population dynamics of marine microbes are strongly influenced by predation from viruses and protists [12, 13]. Far from static, coastal environments exhibit pronounced seasonality, exposing microbes to shifting environmental gradients over time [14]. The combination of well-characterized environmental features and the diversity of single-celled life makes coastal marine environments powerful natural model systems for studying the ecology and evolution of early multicellularity. However, the complexity of this environment also provides a challenge. To move forward, we must develop frameworks that can integrate different ecological drivers of multicellularity.

Here, we ask how the physics of encounters shapes competition for resources among bacteria that degrade organic matter in marine environments. Heterotrophic degraders form the base of microscopic detrital food webs in coastal ecosystems [15–17]. They break down substrates that span at least six orders of magnitude in size — from nanometer-scale organic acids to millimeter-scale marine snow [18]. While interactions between bacteria and large organic particles are known as hotspots for microbial metabolism [19], a much larger fraction of organic matter exists in an intermediate size range [20]: too large and chemically complex for direct uptake by single cells, yet too small to be effectively colonized [21]. We combine experiments and theory to investigate how multicellular group formation influences competition for these intermediate-sized resources. Building on the framework of physical encounter kernels, previously used to model microbial interactions with particles and other cells [22–27], we identify specific regimes where group formation is predicted to increase encounter rates. We also show that a geometric constraint introduces a tradeoff between encounter frequency and per capita resource availability. We explore the ecological consequences of this tradeoff through modeling. Together, our work predicts that the formation of multicellular groups could be a prevalent, but overlooked, strategy for resource competition in marine environments.

## Results

All marine microbes must physically encounter resources in the water column. To build intuition for how cell or group size contributes to resource encounters, we extended an existing encounter kernel framework for marine environments [30, 31] to consider how resources of different sizes are encountered by microbe ranging in diameter from sub-micrometer to hundreds of micrometers. This size range spans from picocyanobacteria to protists, and includes microbes that form aggregates and multicellular groups. The expected rate of geometric encounters between two species *i* and *j* is:

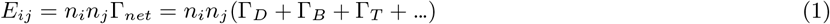

where *n*_*i*_ and *n*_*j*_ are the number concentrations of the individuals *i* and *j*, and the various Γ terms are a summation of encounter kernels driven by different biophysical processes including diffusion (D), buoyancy (B), and turbulence (T). Although encounter kernels for biological processes such as swimming can also be included, initially we omitted these to focus solely on the effects of size. The encounter kernels vary depending on physical factors like the intensity of turbulence and the excess density between the species *i* and *j*. We modeled an energy dissipation of 10^−5^W kg^−1^, which is typical of coastal surf and waves [32, 33], and considered the case where resources and microbes are the same density, with an excess density to seawater of Δ*ρ* = 25kg/m^3^, a value typical of marine microbes and marine snow (see Supplemental section “Varying density”) [34, 35].

This size-dependent view of encounters predicted three regimes (Figure 1a). In the first regime, particulate resources are larger than microbes, and the gradient of the encounter rate is much stronger along the size of the resource, suggesting that resource size is a more important parameter than microbe size. Regime I includes well-studied dynamics such as colonization of marine snow [36–38]. In the second regime, both interacting components are small, meaning that the net encounter kernel is dominated by diffusion. Microbes inhabiting regime II are known to often rely on chemotaxis to forage for ephemeral resource hotspots [24, 39–41]. In the third regime, resources are strictly smaller than microbes, but both pieces are large enough that diffusion is no longer the dominant contributor to the encounter kernel. The microbe’s size is predicted to contribute more to the net encounter kernel in this regime, as evidenced by the horizontal gradient in Figure 1a. Well-characterized strategies of resource acquisition, such as the size-dependent feeding behaviors of ciliates fall in this regime [42]. Notably, regime III is also predicted to encompass a large fraction of interactions among bacterial groups and resources, including interactions spanning particulate and dissolved organic matter.

**Figure 1.**
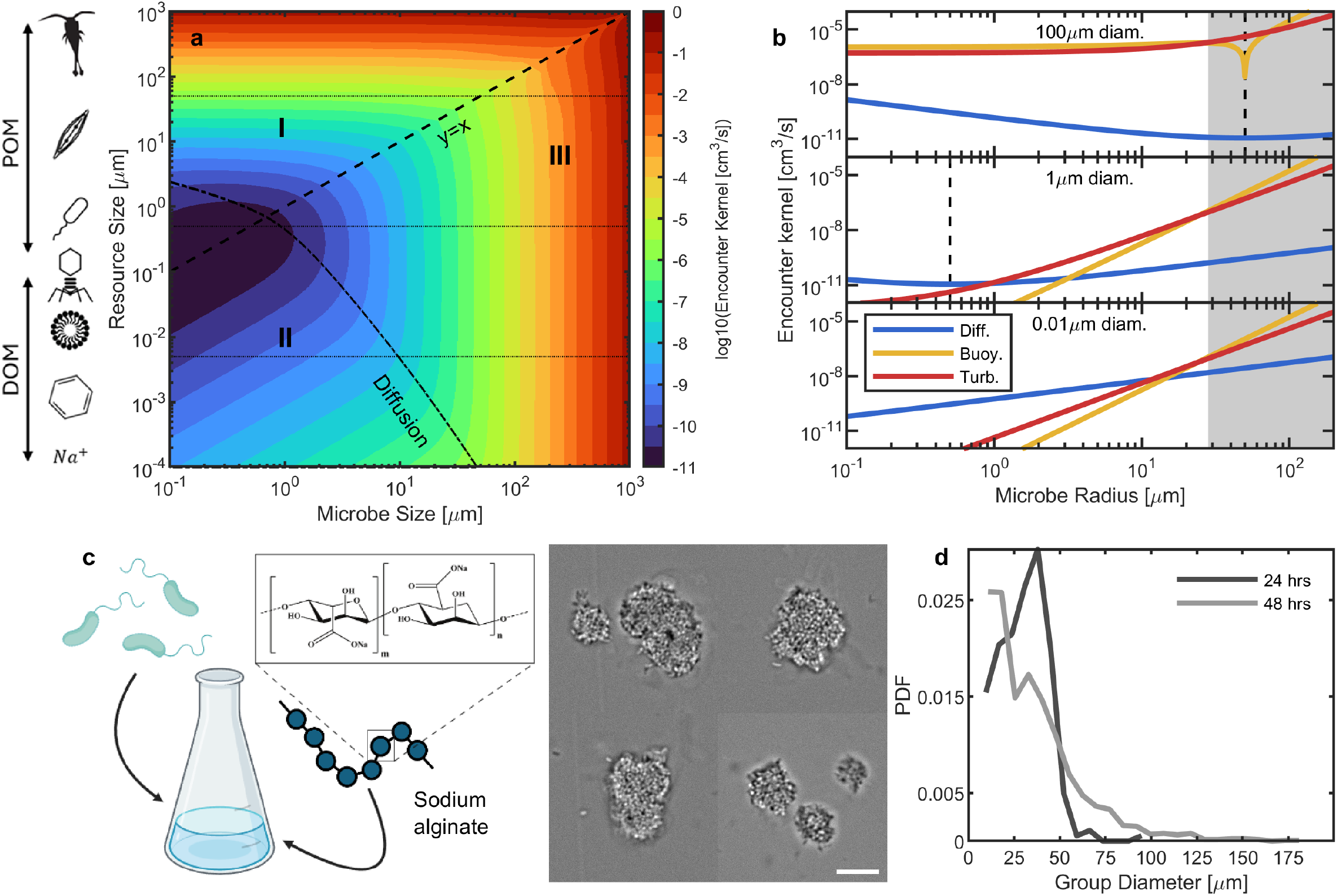
Size-dependent encounters between microbes and resources in coastal oceans predict an advantage for multicellular groups. (a) Predicted size-dependent physical encounter kernel, Γ = Γ_*D*_ + Γ_*B*_ + Γ_*T*_, generated by separately varying the size of resources and the size of microbes. Dashed black line represents *y* = *x*, i.e. where resources and microbes are the same size; dot-dashed line is the contour underneath which diffusion is the dominant contributor to the net kernel. These two contour lines divide the phase space into major sections: we label the 3 largest as Regimes I, II and III. Horizontal dotted lines are cross-sections shown in (b). (b) 1D cross-sections of the encounter kernel in (a) separated into 3 components: diffusion (blue), buoyancy (yellow), and turbulent flow (red). We show theoretical encounter kernels for resources of 100µm diameter (top), 1µm (middle), and 0.01µm (bottom). The gray region indicates the regime where the encounter kernel of turbulent flows is not theoretically resolved [28, 29]. Vertical dashed lines indicate where microbe size is equal to the resource size. (c) When grown on alginate as a particulate carbon source (left), *V. splendidus* strains form multicellular groups (right). Scale bar = 10µm. (d) Size distribution (probability density function, PDF) of *V. splendidus* groups from fluorescence microscopy after 24 hrs (one biological replicate, *N* = 256 groups) and 48 hrs (the same biological replicate, *N* = 1683).

To test the prediction that resource encounters scale with bacterial group size in regime III, we developed a simple experimental model (Figure 1c) based on marine bacteria that degrade the polysaccharide alginate, a linear heteropolymer of uronic acids mannuronate and guluronate found in marine environments. Alginate degradation is a widely conserved trait of the bacterial genus *Vibrio*, and one that drives strain-level diversification [43]. Several strains of *Vibrio splendidus* form multicellular groups when cultured on polymeric alginate [44, 45]. We selected a strain of *V. splendidus*, 12B01, that forms approximately spherical, tightly packed groups of cells when grown on low concentrations of alginate [46]. To measure encounters, we added monodisperse 1µm diameter fluorescent carboxyl-coated microbeads to cultures of bacterial groups. The carboxyl coating of the microbeads was chosen to achieve efficient adsorption to cell groups (2a). Quantification of microbeads attached to bacterial groups revealed the number of microbeads attached per group scaled as *P* = *A*(*r*_*a*_ + *r*_*b*_)^*λ*^ where *r*_*a*_ is the group radius, *r*_*b*_ is the microbead radius, and *A* and *λ* are fit parameters (Figure 2b). Repeating this experiment with varying microbead starting concentrations revealed the prefactor *A* changed with microbead concentration, but the scaling relationship did not (Figure 2c). Such a concentration-independent scaling relationship is consistent with the hypothesis that the primary factor determining microbead acquisition by bacterial groups is physical encounters. Together, these experiments demonstrate that in our experimental model, larger multicellular groups encountered more resources.

**Figure 2.**
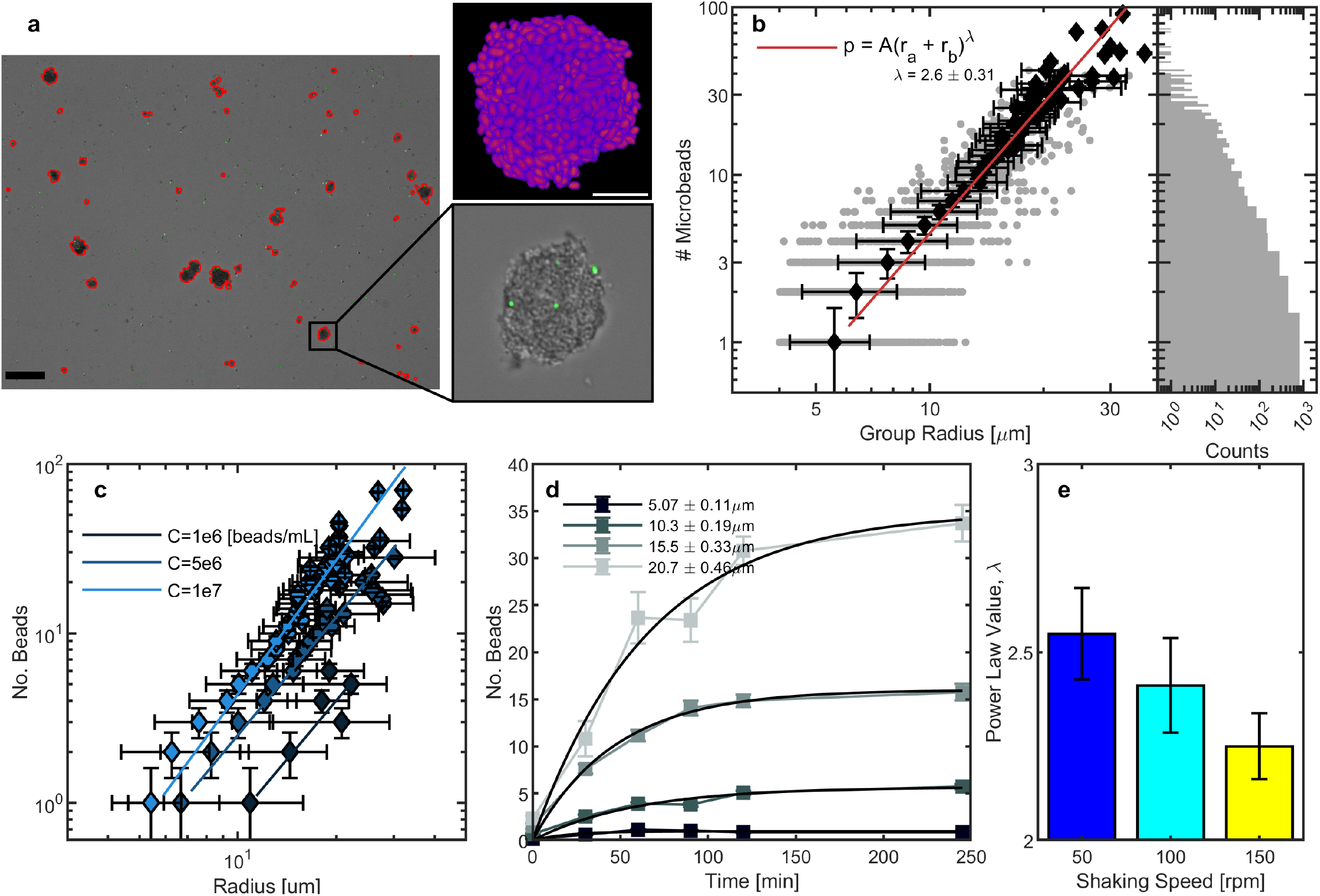
Larger multicellular groups encounter more resources. (a) Example microscope image, showing multicellular groups (DIC), overlaid with the green fluorescent channel showing microbeads (lower inset). Also overlaid in red are the boundaries from image segmentation. Scalebar left is 100µm. Upper inset: A 3D render of a confocal z-stack of one bacterial group, showing dense cell packing. Scalebar is 5µm. (b) Number of microbeads attached vs. group radius for one example treatment shaking at 100rpm for 90 minutes, with *n*_beads_ = 10^7^mL^−1^. The number of groups observed in this single biological replicate was *N* = 3150. Gray points: individual counts of groups with the number of microbeads attached. Black diamonds: average size of groups with a given number of microbeads. Horizontal errorbars are the standard deviation of group radii, vertical errorbars are an estimated error rate from hand-counting (see supplement). Red: power law regression to the mean. Inset: Histogram of the number of groups counted with *n* microbeads. (c) Microbead attachment for different initial concentrations of fluorescent microbeads. Shown is one biological replicate, *N* ={2285, 3130, 3442} for *C* = {10^6^, 5 * 10^6^, 10^7^} beads/mL, respectively. (d) Average number of microbeads attached to groups of different size classes over time. Errorbars are the standard error of the mean for groups in each size class/timepoint. Black lines are fits of a minimal attachment-detachment model (see supplement). Shown is one biological replicate sampled over time, for 10^7^ beads/mL. (e) The value of the power law exponent fit for 3 different shaking speeds tested of one biological replicate. Errorbars are the standard error in the power law exponent from regression.

In addition to physical collisions, we considered the efficiency of microbead attachment and detachment, which could also contribute to the empirically measured size-dependent encounter kernel. We sampled encounters over time for a range of different bacterial group size classes (Figure 2d). In all size classes, the rate at which microbeads accumulated on groups decreased over time, leading to a characteristic plateau of the number of microbeads attached per group. We derived a model of microbead attachment and detachment processes (Supplement Section “Theory of particle attachment and detachment”) that predicted a functional form for this time-dependent relationship that depended on collision rate modified by attachment efficiency, *α*Γ*n*_*b*_, and detachment rate, *β*. We performed a nonlinear regression to the model, finding that the model fit this characteristic curve well (Figure 2d). We found that while particle attachment rate was strongly size-dependent, as expected from our encounter theory, the detachment rate, *β*, was not significantly size-dependent. While further characterization of multicellular group properties is needed to determine the mechanisms through which encounter efficiency changes with group size, the data support the idea that geometric encounters are the primary driver of differences in resource acquisition in our model system.

While the value of the scaling exponent *λ* was empirically determined here, it is worth noting that in some cases, encounter theory predicts a known value for the scaling exponent. For the case where turbulence dominates the encounter kernel, there are simple scaling relationships derived when particles are either much larger or much smaller than the “Kolmogorov” length scale, *η*, which is the size of the smallest eddies in the turbulent fluid. When *r*_*ab*_ = (*r*_*a*_ + *r*_*b*_) ≪ *η*, the predicted scaling relationship is *λ* = 3 [30], and when *r*_*ab*_ ≫ *η, λ* = 7/3 [47]. We might therefore expect that, in the intermediate regime where *r*_*ab*_ ~ *η*, there would be a power law scaling exponent in the range 7/3 *< λ <* 3. This expectation is confirmed in previous numerical studies that explored the finite-inertia regime, which showed that as the particle size increased, the mean encounter rate normalized by the particle volume decreased [29]. We found that the measured scaling exponent *λ* fit directly within the expected range from encounter theory. In real ocean surface systems, it is predicted that the Kolmogorov length will generally be larger than about 100 µm, and will generally be larger than 1 mm below about 10m depth [32, 48]. We cannot theoretically define the Kolmogorov length scale in our shaken culturing flasks, but we reasoned that if the Kolmogorov length scale contributed to the net encounter kernel scaling exponent, altering the shaking speed should change the net encounter kernel exponent. This is because the Kolmogorov length scale is inversely related to the intensity of turbulence in a fluid [47], which can be modified by changing the shaking speed. In our experiments, as shaking speed decreased, the empirically measured value of *λ* increased (Figure 2e), indicating that turbulence plays a strong role in setting encounters in our system.

Having characterized the “mean-field” behavior of the net encounter rate, we next explored a defining feature of patchy, heterogeneous systems: fluctuations around the mean. We hypothesized that the distribution of encounters would be Poisson (that is, the probability density function for k encounters is *f* (*k*) = *γ*^*k*^*e*^−*γ*^/*k*! for each size class), which is expected in situations where encounters occur independently with some known mean rate *γ*. A Poisson prediction for the distribution of encounters in each size bin matched the data well for small groups (Figure 3a and Supplemental Figure S8). However, the mean and variance diverged when multicellular groups reached larger group sizes (Figure 3b), starting around 15µm and larger (Figure 3c). We found that a size-dependent deviation from Poisson predictions was consistent across different shaking speeds, flask sizes, and shaking modalities, although the exact size at which the mean and variance diverged varied in a condition-dependent manner (see Supplemental Figure S9). Contributing to this, we found a negative correlation between group size and group sphericity (Supplemental Figure S10). Filtering aspherical groups from our dataset led to stronger agreement between the mean and variance in the number of microbeads attached across size bins. Together, these results suggest that microbeads on small multicellular groups are acquired in a Poisson manner, while larger multicellular groups may have distinct properties, such as aspherical shapes, that prompt deviations from the Poisson distribution.

**Figure 3.**
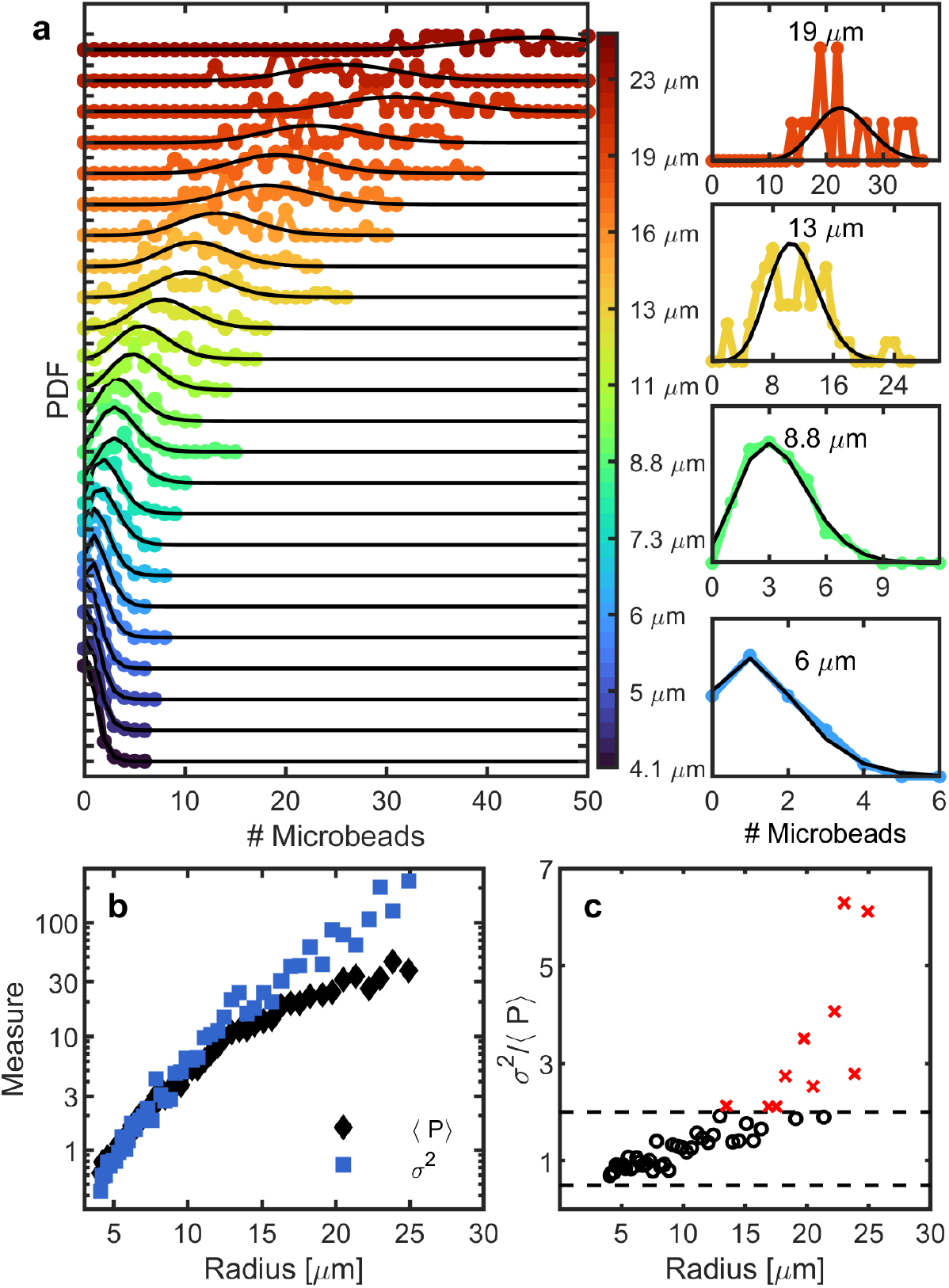
Encounters between resources and individual groups are Poisson-distributed. (a) Histograms of the number of microbeads attached to multicellular groups for the same biological replicate as shown in Figure 2 at 120 minutes of incubation with microbeads. Each color histogram represents a different size class and is offset for visualization. Black lines represent Poisson predictions given the same mean as the size class. Thicker lines are size classes shown to the right. PDF is the probability density function. Right, selected, representative histograms of the number of microbeads attached on multicellular groups for 4 different size classes. (b) Average (black diamonds, ⟨*P*⟩) and variance (blue squares, *σ*^2^) in the number of microbeads attached vs. group size, plotted on the same y-scale. These values are extracted from the data in Figure 3a (c) Variance divided by mean for each size bin (*σ*^2^/ ⟨*P*⟩). Horizontal dashed lines are min and max bounds of the chosen test statistic, at 0.5 ≤ *s* ≤ 2 where *s* = *σ*^2^/Γ. Points inside these bounds are displayed as black circles, outside as red x’s.

How might a positive relationship between size and encounter rate shape the ecological dynamics of group-forming bacteria? To understand this question, we examined the *per capita* distribution of resources on multicellular groups. We first estimated the number of cells in multicellular groups of different sizes by enumerating the number of cells in 10 multicellular groups using confocal microscopy (see methods). We found that the volume of multicellular groups scaled linearly with the number of cells in the multicellular groups, allowing us to consistently convert apparent size via transmission microscopy to number of cells (Supplemental Figure S11). We then converted our data (Figure 2b) to a *per capita* distribution of resources, and found that while the mean number of microbeads per cell decreased with group size, so did the variance in the number of microbeads per cell (Figure 4a). Put another way, smaller groups had a higher mean rate of resource encounters per capita, but there was also high variation in whether any one group encountered resources. Many small groups encountered no resources over the measured time interval. Therefore, increasing size resulted in fewer, but more consistent, resource encounters per capita (Figure 4b). Intriguingly, the relationship between consistency and mean resource per capita appears concave (Figure 4c), suggesting a tradeoff that maximizes either of the extremes - larger cellular groups have high consistency to encounter low amounts of resource, and smaller cellular groups have higher potential to strike it big on the resource lottery.

**Figure 4.**
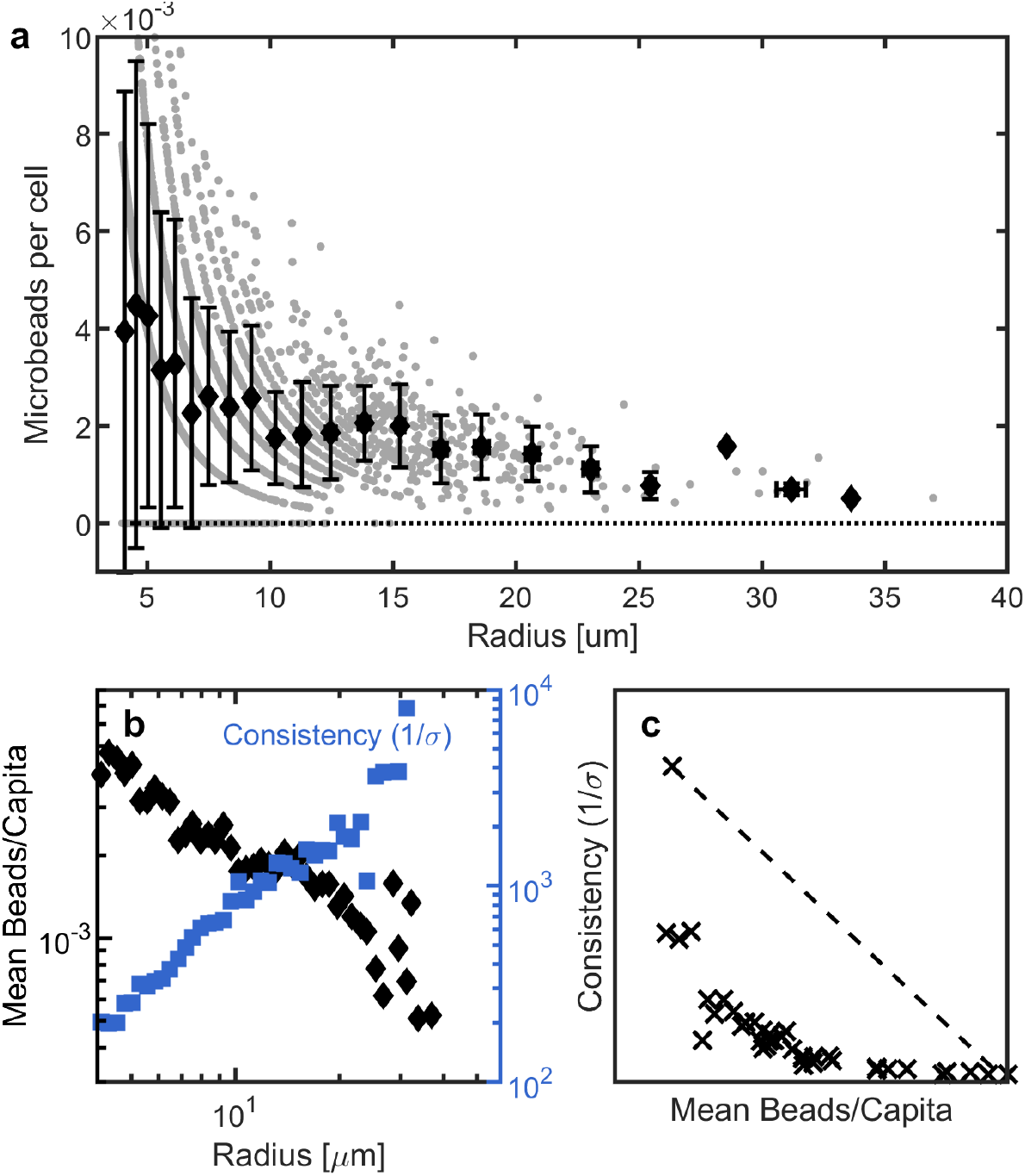
Per capita resources encountered suggest a size-dependent trade-off in the mean rate and the consistency of encounters. (a) Number of microbeads attached to a multicellular group per number of cells in the group. Gray points: individual instances, black diamonds: averages for different multicellular group size classes. Every 5th radius bin is shown for clarity. Horizontal black line represents encountering 0 microbeads per capita. Data is replotted from Figure 2a. (b) Mean and consistency (inverse of the standard deviation) of microbeads attached per capita from (a). (c) Consistency plotted vs. the mean number of microbeads per capita. Dashed black line represents neutral selection, i.e. no directional advantage.

To understand the consequences of a tradeoff between consistency and per capita resource encounters for the dynamics of resource competition, we developed a simple computer model to simulate the essential features of our experimental system: namely, fluctuations in resource encounters, a rate of encounters that is size-dependent, and the ability to grow as either individuals or multicellular groups (Figure 5a). In this model, populations can exist as a mix of single cells (which we called state S1) and multicellular groups (state S2). In state S1, cells divide and separate after division. In S2, cells divide but do not separate, instead remaining in the group and increasing group size. The growth of both states is dependent on resource encounters via Monod kinetics. We simulated populations of single cells that were initially proportioned into equal halves occupying either the S1 or S2 state.

**Figure 5.**
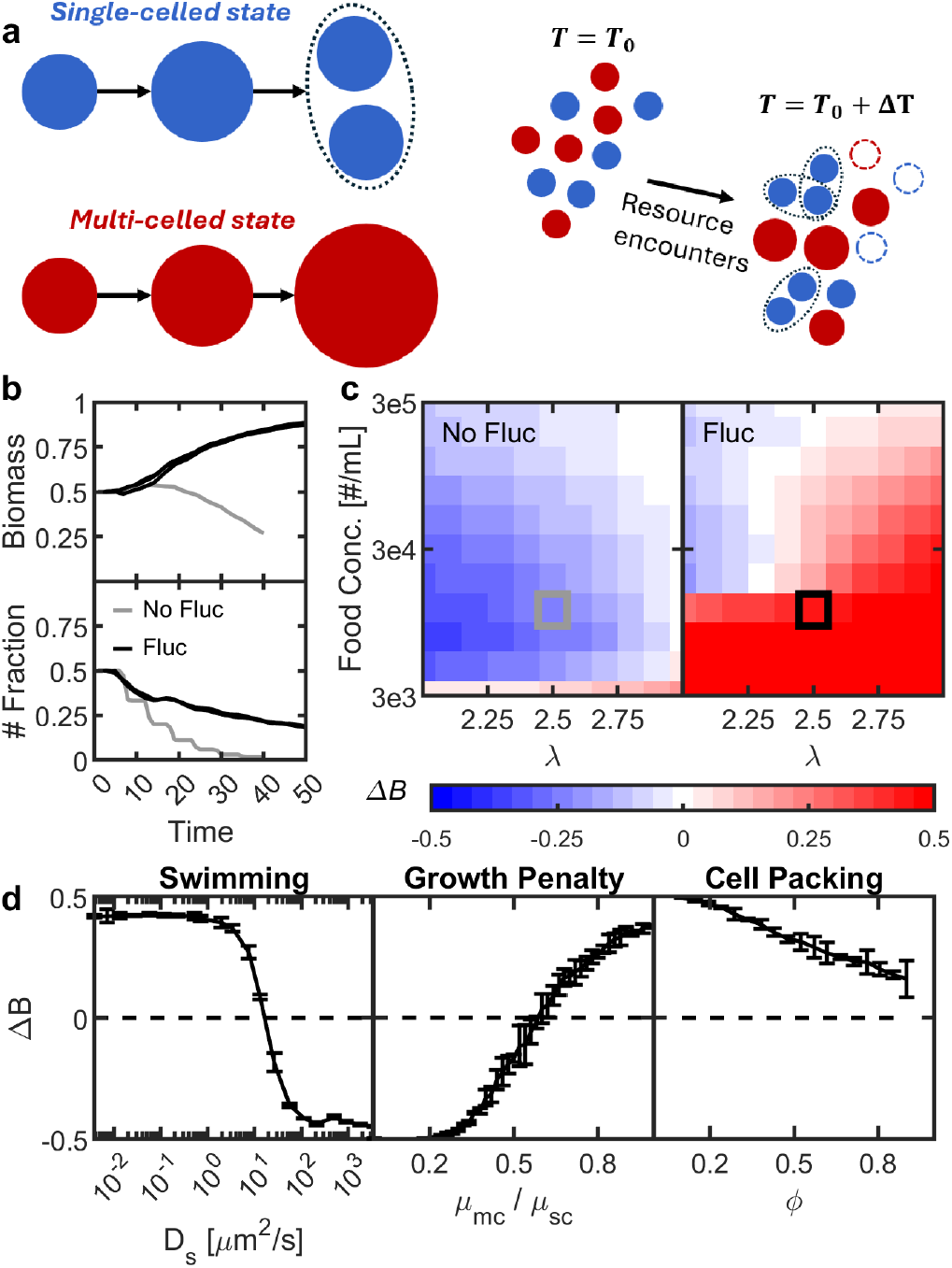
A model of encounter-driven resource competition predicts trade-offs between the mean-rate and the consistency of resource encounters define regimes where multicellular groups are the more competitive growth strategy. (a) Two different cell states in the model. The single-cell state divides when it reaches double its original size, the multi-celled state does not divide, instead simply gaining biomass per individual. (Right) An illustration of how the simulation works in practice, illustrating division (dashed black outlines), growth (increase in size) and death (dashed outlines) as processes based on resource encounters. (b) Time tracks of the number fraction (bottom) and biomass fraction (top) of the multicellular state, for *N* = 3 simulations for each type. Gray: simulation in only the mean field, i.e. not allowing for fluctuations in the number of encounters. Black: Tracking the multicellular state in a fluctuating, patchy environment. (c) A heatmap of the average change in biomass fraction of the multicellular state without (left) and with (right) fluctuations in the number of resources encountered, swept over the concentration of food and different values of *λ*, the net encounter kernel scaling exponent. Simulations from (b) are boxed. *N* = 3 simulations for each parameter combination. (d) Change in biomass fraction for the multicellular state when including swimming for the single-celled state, growth penalties, and different cell packings. For each parameter set, *N* = 3 simulations were averaged. Shown is 2*σ* = 95% confidence bounds. Resource concentration was chosen such that the mean rate of encounters was *G*_0_ = 1.2 * 10^−1^ encounters per fastest doubling time for the single-celled state, and *λ* = 2.7.

Our primary goal was to compare mean-field and stochastic models to understand how fluctuations in encounters change the ability for single-celled and multi-celled states to compete for resources. We swept two variables: the concentration of food particles *c*, and the scaling exponent of resource acquisition, *λ*. A different value for the scaling exponent *λ* could be achieved by either a different environment (for example, a change in turbulence levels) or by a variable size-dependent attachment efficiency or detachment rate. When there were no fluctuations in resources (gray lines, Figure 5b), the total biomass of S2 and the number of S2 groups both decreased due to resource out-competition by the S1 state. We found that S1 out-competed S2 in the mean field simulation over a wide range of resource concentrations and encounter scaling values (Figure 5c). At very low resource concentrations (Supplemental Figure S11), both S1 and S2 states died off. Regimes of coexistence emerged as lambda approached 3, corresponding to conditions where there is less turbulence and/or more efficient resource encounters at larger sizes.

Next, we introduced fluctuations in resource acquisition via Poisson sampling. In our simulations of fluctuating resource encounters (black lines in Figure 5b), we found broad regimes in which S2 multicellular states out-competed S1 in terms of total biomass (Figure 5c). These regimes emerged even as the total number fraction of S2 entities decreased, as in our model they have no mechanism for replicating. Notably, S1 could only outcompete S2 when resource concentrations were high and *λ* was low, corresponding to inefficient resource encounters or highly turbulent environments. Collectively, these simulations predict that the ability to form large groups, and experience consistent, but low per capita resources, can present an advantage for resource competition in many types of fluctuating environments.

To further understand how the properties of single-celled and multi-celled states interact with stochastic encounters, we asked whether swimming in the S1 state could tip the competitive balance. We increased the encounter rate of S1 to emulate swimming by adding a non-chemotactic swimming term to the resource acquisition encounter kernel (see Methods). The strength of swimming is quantified by the effective diffusion coefficient, *D*_*s*_, which we allowed to range from 3 * 10^−3^ to 3 * 10^3^µm^2^/s. This corresponds to swimming velocities from 0.1 to 100µm/s, when bacteria exhibit run-and-tumble motion reorienting at a rate of 1Hz. If bacteria tumble more frequently, this range corresponds to higher swimming speeds (see Supplemental Figure S12). Recovering mean-field behavior (i.e., recovering a regime where S1 outcompetes S2) required a diffusive capacity of 17µm^2^/s, corresponding to swimming speeds of 7µm/s when reorientations occur once a second, and *v*_0_ = 50µm/s when reorientations occur at a rate of 10Hz (Figure 5d).

We also tested two other types of scenarios that could weight a competition more towards the single-celled S1 strategy: density-dependent growth penalties, and varied density of cell packing (Figure 5d). We reasoned that both might affect growth and resource distribution in the multicellular S2 state, as local density can increase resource competition, and this effect could be exacerbated if cells are packed more densely. To add each effect to our model, we imposed a growth penalty, quantified as the ratio of the maximum growth rates between the two states, *µ*_*mc*_/*µ*_*sc*_, and included packing fraction, *ϕ*, which shifted the conversion for the number of cells in a group to the size of that group. A growth penalty changes per capita survival, while cell packing changes the per-capita resource encounter rate. We tested the same food concentration and size-scaling exponent as the swimming case above. We found that the S2 state required a growth penalty of about 60% of the S1 state in order to destabilize regimes in which coexistence had been observed (Figure 5d). However, even cell packing up to 90% of the total volume of the multicellular group could not destabilize coexistence. As a 90% packing density corresponds to a very dense packing, these simulations suggest that competition in our simulation is sensitive to growth penalties, but not cell packing.

## Discussion

In coastal marine environments, microbial ecology is shaped by physical encounters. Here, we highlight size dependence as an important feature of encounters (Figure 1). In competitive settings, we predict a regime where forming multicellular groups—and thus increasing size—enhances a cell’s ability to encounter resources (Figure 5). We propose this as a resource acquisition strategy that contrasts with motility-based foraging, such as swimming and chemotaxis, which have been widely explored in the optimal foraging literature [49]. Notably, the swimming speed predicted in our model of competition among single cells vs. multicellular groups to shift the growth advantage of multicellular groups is generally approachable by marine bacteria [50, 51]. But while motility increases encounter rates by expanding the search area, it comes at a metabolic cost. The tradeoff between expending energy and increasing encounters defines timescales over which cells can remain motile [52–55], limiting how long cells can remain active. In contrast, group formation may sustain resource acquisition over longer timescales without the energetic costs of motility.

A prediction from our work is that, *per capita*, cells in multicellular groups encounter fewer resources that single cells (Figure 4). In practice, this effect could be offset by processes that depend positively on cell density; for instance, growth on polysaccharides often require interactions among cooperating cells to pool extracellular enzymes required for catabolism of these resources [45, 56]; alternatively, cross-feeding interactions can allow some cells to proliferate on the byproducts created by the metabolism of degraders, forming a microbial food web [57, 58]. In environments where resources are sparse, both crossfeeding and cooperation will depend on microscale spatial structure to create local density [44], or a ‘neighborhood’ of interaction. Our resource competition model suggests that the architecture of multicellular groups, measured through their packing fraction, can offset growth penalties associated with the geometric constraint of resource encounter (Figure 5). Changes to the structure of multicellular groups can be achieved by altering cellular traits such as shape and surface properties [59–61]. A limitation of the present work is that we do not consider within-group allocation of resources. However, by expanding the model that we present in the current study, it will be possible in future work to address how the metabolism and physical structure of groups overcomes the geometric constraints of group size.

Our empirical measurements reveal an important aspect of marine environments: resource encounters are stochastic (Figure 4). We predict that, in the context of resource competition, the tradeoff between the amount and consistency of resource encounter defines broad regimes where multicellular groups are the most competitive ‘strategy’ of resource competition. This idea is congruent with Tilman’s resource competition theory [62], which predicts that when resources are scarce, the most fit competitor will the the one that minimizes population level attrition through more efficient growth and enhanced survival. Our findings raise the possibility that, in many marine environments, consistent resource encounters by multicellular groups may be a key strategy to survive resource scarcity.

While we focus on resource encounter in this study, resources are just one type of type of particle that can be encountered. The spectrum of physical encounters encompasses predators [63], phage [13], potential partners to horizontally transfer genetic information, or competing cells competition [44, 58, 64, 65]. In future work, our encounter-based framework can readily be extended to encompass these other types of physical interaction to connect the physical processes of encounters to their ecological and evolutionary consequences.

## Materials & Methods

### Data and Code

All scripts used for image analysis, simulations, and plotting figures can be found here: (https://github.com/thomas-c-day/project-encounter.git).

### Bacterial cultures and growth conditions

*Vibrio splendidus* strain 12B01 [66] was routinely cultured in Marine Broth (Difco 2216), with 1.5% agar for solid medium. Cultivation was carried out at 25 °C. Cultures of 12B01 grown as a multicellular groups were established as described previously [46]. Briefly, cells were grown as a preculture in 10mM glucose MBL minimal medium to an optical density (OD) measured at 600 nm of 0.2-0.5. OD measurements were taken using a Genesys 20 Spectrophotometer (Thermo Scientific). To prepare inoculum for cultures, cells were pelleted at 6000 rcf for 1 min in a tabletop microcentrifuge (Eppendorf), and resuspended in carbon-free MBL minimal medium at a final OD of 1.0. Experimental cultures were initiated by inoculating 12B01 at an initial density of 10^−5^ OD into 1.5mL of MBL minimal medium containing 0.07% w/v low viscosity alginate (Sigma A1112) in a 24 well-plate culture vessel (VWR 10861-558). Cultures were grown with shaking at 125rpm on a Thermo Scientific MaxQ 2000 orbital shaker, unless otherwise specified.

### Microbead incubation experiments

Bacterial groups were grown in liquid cultures overnight, such that cultures contained a range of bacterial group sizes. One micron diameter carboxyl-coated fluorescent microbeads (Bangs Laboratories) were sonicated for 15 min at 40 KHz (Sonicator: Branson 5800) to minimize microbead-microbead adhesions. We then mixed bacterial groups from culture and microbead suspensions at various relative concentrations. For the data shown in the main text, we used microbead concentrations of *c*_*b*_ = {10^6^, 5 * 10^6^, 10^7^} beads per milliliter, measured by diluting the suspensions from the manufacturer’s stated bead concentrations.

Incubation occurred in 1 well of a 24 well plate, filled with 1 mL of fluid for up to 245 min. The cultures of bacterial groups and microbeads were set on an orbital shaking platform at various speeds, as indicated. At each sampling point, we removed the culture from the shaking incubator and sampled 30 µm from the mixture onto a polymer-bottom well slide (Ibidi), then imaged. For experiments where we changed shaking speed, we separately varied the orbital shaking speed to be 50, 100, and 150rpm for three separate cultures. For experiments where we changed the shaking type and container, we used three separate setups described here: first, we used the typical setup described above - 1 mL culture with beads in a 24 well plate, shaken on the same orbital platform at 100 rpm. Second, we used a much larger volume of fluid - 15 mL of bacterial culture contained in a 25 mL glass culture flask - set on the same orbital shaker at 100 rpm. Third, we used the typical volume and plate setup with a different shaker, a side-to-side shaker set at 107rpm (IKA-Werke HS501).

### Microbead microscopy

Images were taken with a Nikon Eclipse Ti2-E inverted microscope with 10x Nikon Plan Apo DIC air objective (NA 0.45) and an additional 1.5x zoom, for a net magnification of 15x, with a Orca Fusion C14440 digital camera and LED lightsource (Nikon D-LEDI). To ensure that all microbeads attached to the cell groups were visualized, we imaged multiple z-planes spaced at 5 µm intervals in both transmission DIC mode and in the green fluorescent channel (fluorescent excitation 488 nm). Each z-stack was then projected into a single image using the Nikon Elements software (AR 6.02.03) “Extended Depth of Focus” function, a method of obtaining a z-projection of maximum focus. The DIC (transmission) channel was segmented using a trained model of Ilastik [67] and custom-written MATLAB scripts (see Supplemental material for scripts) to generate the boundary of each aggregate larger than 4 µm in diameter. The fluorescent channel was segmented in MatLab (version R2024a) to find microbead centers, and the number of microbeads that resided within each aggregate boundary were enumerated.

In order to ensure that microbeads were not counted as a very small cell group with a microbead attached, we measured the total fluorescence per area of each group identified. If the value of the fluorescence per area exceeded 5 * 10^3^ units, we discarded the measurement.

### Measuring number of cells per group

Cellular packing within individual groups was measured for 10 bacterial groups via confocal microscopy. Cell stain Syto-9 (Invitrogen) at 5 uM concentration was used to label individual cells of *V. splendidus* 12B01 grown on alginate for 36 hrs. We used a point-scanning confocal microscope (microscope body: Lecia DMI4000B, confocal unit: Leica TCS SPE) equipped with a solid-state 488 nm laser unit, and 2 photomultiplier tubes with an ACS APO 40x/1.15 NA CS series oil objective to image cells within groups with pixel resolutions between 0.135µm and 0.144µm per pixel. Z-stacks were taken with z=0.40 µm, and the photomultiplier gain was set separately for each stack to maximize the dynamic range without saturation. After imaging, the xy and z resolutions were made equal using Fiji [68] release 2.15.0 and its plugin “Make isotropic”. Then, we used a custom Matlab script to segment the images, which included binarization, watershedding, and ellipsoid fitting.

### Cell state competition simulations

We developed a simulation to model competition between two microbial strategies - single-celled and multicellular states — each drawing resources from a common pool. Individuals grow and decay based on the amount of food they acquire, like *db*/*dt* = (*µ* − *δ*)*b*, where *b* is the current total biomass of the individual, *δ* is a constant decay rate that penalizes starvation, and *µ* is a Monod-like growth rate that varies based on the per-capita food encountered. In the single-celled state, individuals divide once their biomass doubles; in the multicellular state, they do not divide and continue growing. Death occurs when biomass drops below a critical threshold. The simulation proceeds in rounds, with each individual’s biomass updated based on net growth, division, or death.

Food encounters were computed using empirically motivated encounter kernels incorporating diffusion, turbulence, and, in some cases, swimming. Swimming was modeled as run-and-tumble motion, contributing an effective diffusion term to encounter rates. We simulated both deterministic and stochastic (Poisson-distributed) food availability, and explored a range of swimming strengths. To assess trade-offs, we varied the maximum growth rate of the multicellular state to introduce penalties relative to single-celled growth. Full simulation details, including equations and parameter sweeps, are provided in the Supplementary Methods.

## Acknowledgements

We thank Jonasz Slomka for helpful discussions about the physical encounter kernel. T.C.D was supported by the Life Sciences Research Foundation and by a grant from the Simons Foundation (Day-24-0801-A0001). J.S. acknowledges startup support from the University of Southern California.

