## Supplemental figures and methods for "Multicellular group formation as a resource competition strategy for marine bacteria"

#### Theory of encounter rates

The rate of encounters between two particle species $i$ and $j$ can be modeled via:

$$E_{ij} = \Gamma_{ij} n_i n_j \tag{1}$$

where $n_i$ is the concentration of species $i$ in particles per unit volume. The encounter kernel $\Gamma_{ij}$ has units of volume per time, meaning if a particle of type $i$ (such as a group of bacteria cells) can "see" more fluid per unit time, then it will encounter more particles of species $j$ (for example, a particle of food). The encounter kernel is affected by a variety of different kinds of biophysical processes, including particle diffusion, advection of the particle embedded in a fluid, buoyancy differences, and active motility. In turn, each of these terms of the net encounter kernel is affected by the particle sizes; we write this here like so:

$$\Gamma_{ij} = \Gamma_D(r_i, r_j) + \Gamma_B(r_i, r_j) + \Gamma_T(r_i, r_j) + ... \tag{2}$$

where $r_i$ and $r_j$ are the radii of the particles under consideration. The relative effects of these processes on the encounter kernel have been detailed elsewhere [1, 2]; here, we reproduce the simplest versions of these processes for comparison between them.

The diffusion encounter kernel is:

$$\Gamma_D = 4\pi(D_i + D_j)(r_i + r_j) \tag{3}$$

where $D_i$ and $D_j$ represent the thermal diffusion constants.

The buoyancy kernel is:

$$\Gamma_B = \pi(r_i + r_j)^2 |u_i - u_j| \tag{4}$$

where $u_i$ represents the vertical velocity of particle $i$, and depends on particle density and size.

The turbulence kernel is:

$$\Gamma_T = 1.3(r_i + r_j)^3 \sqrt{\frac{\epsilon}{\nu}} \tag{5}$$

when the objects are much smaller than the Kolmogorov length scale. Here, $\epsilon$ represents the energy dissipation rate, and $\nu$ represents the kinematic viscosity of the fluid. The Kolmogorov length scale is also related to these parameters like $\eta = (\epsilon/\nu)^{1/4}$. When much larger than the Kolomogorov length scale, the turbulence kernel is instead [3]:

$$\Gamma_T = 1.37\pi\epsilon^{1/3}(r_i + r_j).^{7/3} \tag{6}$$

Note that these two functional forms do not intersect at the Kolmogorov length scale.

For ballistic swimming (i.e. swimming in a straight line), the encounter kernel is:

$$\Gamma_S = \frac{4}{3}\pi(r_i + r_j)^2 u \tag{7}$$

where $u$ represents the swimming velocity of a motile cell in relation to a stationary particle. By inspection, we can see that this kernel scales quadratically in the size of the particles, but only linearly in the swimming speed. In this sense, all else being equal, changing size is a more important variable than changing swimming speed on the rate of encounters. Of course, microbes do not generally swim in a straight line. Often, they behave in a "run and tumble" fashion, where runs of roughly ballistic swimming are punctuated by brief periods of reorientation. The resulting trajectory is classically modeled

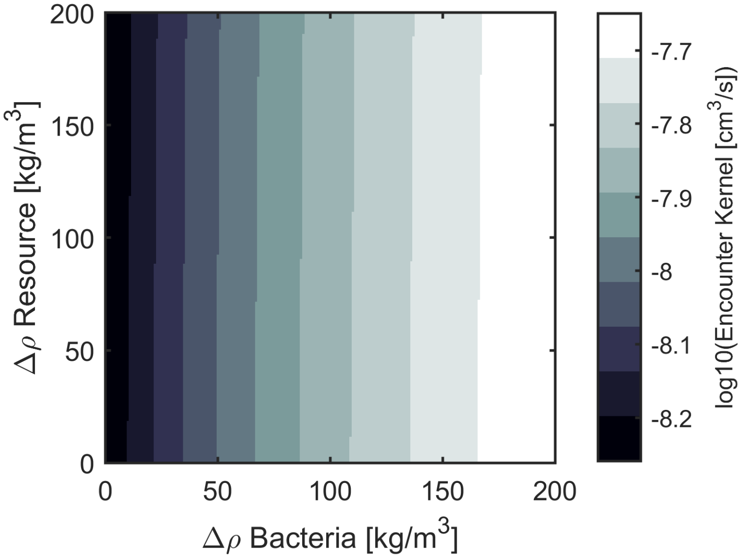


**Figure S1: Net encounter kernel varies with changing particle densities**. The net encounter kernel as a function of excess density ( $\rho$ ) of 10 µm bacterial aggregates and 1 µm resource patches.

as a random walk, at least in the case where there is no chemotaxis. Random walks have the same characteristics of thermal diffusion, meaning that we can use the same form of the diffusion encounter kernel here, but with the thermal diffusion coefficients instead replaced by effective diffusion constants set by the mean squared displacement of bacterial swimming. The mean squared displacement is, in turn, set by run speed and run lengths. In later simulations of encounters via cell swimming, we will therefore approximate the encounter kernel like $\Gamma_S = 4\pi D_i(r_i + r_j)$ where $D_i$ represents the effective diffusion constant of swimming, and is determined by the mean squared displacement of the random walk of bacterial run-and-tumble motion like: $\langle (x - \bar{x})^2 \rangle = 6D_i t$. When there is chemotaxis in swimming, the random walk formulation of swimming is not as appropriate.

#### Varying the density of particles

Differences in buoyancy are an important driver of encounters in the ocean. The buoyancy encounter kernel is:

$$\Gamma_B = \pi(r_i + r_j)^2 |u_i - u_j| \tag{8}$$

where $r_i$ and $r_j$ are the sizes of the interacting species, and $u_i$ and $u_j$ refer to their velocities in the water column, which can be either positive (for sinking particles) or negative (for rising particles). The velocity $u_i$ is generally taken to be the terminal velocity of a (floating or sinking) object, and depends on particle density and shape. In the case of a small, spherical, sinking particle at low Reynold's number, the terminal velocity is $|u_i| = 2\Delta\rho g r.^2/9\nu$, where $\nu$ is the dynamic viscosity, $g$ is the gravitational constant, $r$ is the particle size, and $\Delta\rho$ is the difference in density between the particle and the water. Particles that are 10 µm in radius and 10 g/cm$^3$ in excess density of water approach their terminal velocity in less than ten-thousandths of a second, so the terminal velocity is quite a good approximation.

There is a broad range of particle densities in the ocean. Marine snow has been measured to have excess densities ranging from $10^{-5}$ to $10^{-1}$ g/mL [4], and similar values have been measured for living plankton [5]. For the main text Figure 1, we chose the middling value of $\Delta\rho = 0.025$ g/mL $= 25$ kg/m$^3$, and considered the case where microbial aggregates and resources shared the same excess density. We were interested in extending our analysis to consider situations where bacterial aggregates and resources do not share the same excess density. In Figure S1 we sweep a range of excess densities, from $0 \leq \Delta\rho \leq 200$ g/mL, separately for a bacterial aggregate 10 microns in radius and a resource patch 1 micron in radius, and calculate the expected net encounter kernel when $\epsilon = 10^{-5}$ W/kg. The encounter kernel varies by less than an order of magnitude from the extreme ends of the parameter sweep. We also found that the larger particle contributes more strongly to the gradient in the encounter kernel.

### Theory of particle attachment and detachment

By observing the average number of beads per multicellular group after a certain time interval, we are observing the encounter rate modified by both an attachment efficiency and some detachment process. We write:

$$\frac{dp}{dt} = \alpha\Omega - \beta p \tag{9}$$

where $p$ enumerates the average number of beads attached onto one multicellular group, $\alpha$ is the attachment efficiency of beads to multicellular aggregates, $\Omega = n_p\Gamma$ is the encounter rate of a multicellular group with any bead, $n_p$ being the concentration of beads in the solution, and $\Gamma$ representing the size-dependent geometric encounter kernel, and $\beta$ is a detachment rate of beads from groups. We assume that $\alpha$ and $\beta$ are independent of how many beads are attached to the group, and are independent of time. In general, both $\alpha$ and $\beta$ can be size-dependent.

#### Particle attachment without any bead depletion effects

First, we consider the limit of high bead concentration, *i.e.* we do not consider dynamic changes to the concentration of beads that are not attached to any multicellular groups. We will write this as $n_p(t) = n_0$. Further, we assume that the multicellular group concentration changes negligibly in time.

The above equation is a linear differential equation of the form

$$\dot{p} + f(t)p = g(t) \tag{10}$$

which is exactly solvable. The solution is

$$p(t) = \frac{\alpha}{\beta}\Gamma n_0 + Ke^{-\beta t} \tag{11}$$

With the initial condition that $p(0) = 0$, we solve for the integration constant $K$, so that the final time-dependent result is

$$p(t) = \frac{\alpha}{\beta}\Gamma n_0(1 - e^{-\beta t}) \tag{12}$$

which can be confirmed as a solution to the differential equation above by substitution.

The variables $\alpha$ and $\Gamma$, representing the attachment efficiency and the encounter rate, appear together in this equation. Therefore, the attachment efficiency is not possible to measure from timetracks of the number of beads attached. We therefore combine the variables $\alpha$ and $\Gamma$ in Equation 12 into one term (which we will call simply $\Gamma$), which represents the geometric encounter kernel modified by the attachment efficiency. There are therefore two free parameters in this equation: $\Gamma$ and $\beta$.

#### Including the effect of microbead depletion during incubations

In the case where microbeads are constantly being depleted from the culture (for instance, because there is no influx of new microbeads), then the situation derived above changes to include information about the time-dependent microbead concentration. Here, we will show that the form of the solution to this situation is not dramatically different than that above, and approaches the same form as above in the limit of low concentration of multicellular groups.

We model the concentration of microbeads in the culture $n(t)$, which is a function of time as they get depleted. The concentration of microbeads attached to multicellular groups is $a = pc$, where $p$ is the number of microbeads attached per multicellular group, and $c$ is the concentration of multicellular groups. The total concentration of beads $N = n + a$ remains constant, *i.e.* microbeads either must exist in the culture medium or attached to multicellular groups. We can write this like:

$$0 = \frac{dN}{dt} = \frac{dn}{dt} + c\frac{dp}{dt} \tag{13}$$

The number of beads attached per multicellular group can be written, like above:

$$\frac{dp}{dt} = \alpha\Omega - \beta p \tag{14}$$

$$= \alpha\Gamma n - \beta p \tag{15}$$

Differentiate both sides to obtain

$$\ddot{p} = \alpha\Gamma\dot{n} - \beta\dot{p} \tag{16}$$

where we use dot notation for brevity. Substituting for $\dot{n}$ using Equation 13,

$$\ddot{p} = -(\alpha\Gamma c + \beta)\dot{p} \tag{17}$$

We solve this equation for $\dot{p}$, with the initial condition that $\dot{p}(0) = \alpha\Gamma N$.

$$\dot{p}(t) = \alpha\Gamma N e^{-(\alpha\Gamma c+\beta)t} \tag{18}$$

which we can integrate

$$p(t) = \int_0^t \dot{p}(t')dt' \tag{19}$$

$$= \frac{\alpha\Gamma N}{\alpha\Gamma c + \beta}(1 - e^{-(\alpha\Gamma c+\beta)t}) \tag{20}$$

In the limit where $\beta \gg \alpha\Gamma c$, then this approaches the solution of Equation 12.

We next sought to understand if it would be more valuable to include the depletion effect in our models. Using the parameters $\beta = 0.01\ \text{min}^{-1}$, and $\Gamma = 1 * 10^{-8}\text{mL}\,\text{min}^{-1}$, with 100% attachment efficiency, this would require a concentration of multicellular groups to be greater than 1 million groups per milliliter for the ratio $\alpha\Gamma c/\beta > 0.10$. If an OD 1 suspension of bacterial cells all clumped into groups with 1000 cells, then we expect the concentration of multicellular groups in suspension is likely to be at least one order of magnitude below this value. We therefore concluded that the non-depleting solution was adequate to model our dataset.

#### Fitting the attachment-detachment model to microbead experiments

In our experimental data, we noted that the distribution of microbead attachment to larger multicellular groups was less well fit to our empirically measured encounter kernel. We sought to understand the main contributor to this size-dependent behavior of microbead attachments. To do this, we first considered the scenario where both $\Gamma$ and $\beta$ may be size-dependent. We used a nonlinear least squares regression to fit the model Equation 12 to experimental data via the following method. First, we binned timelapse data into distinct size bins (by group radius). We excluded size bins that had fewer than 10 instances. In each bin, the number of microbeads per multicellular group was averaged to obtain a series of timetracks, which we could fit with our model. We chose an initial parameter guess to be $\beta = 0.01\text{min}^{-1}$, and $\Gamma = 1 * 10^{-8}\text{mL}\,\text{min}^{-1}$ for each timetrack. We used MatLab's nonlinear fitting algorithm fitnlm. The respective fits for a few select size bins are summarized in Table S1, and the fits along with the time-tracks for various size bins are displayed in Figure S2a.

This fitting procedure found strong size-dependence for the fitting parameter $\Gamma$, representing the particle attachment rate, as expected from encounter theory. The Pearson's r-correlation coefficient was 0.97, and a Mann-Kendall trend test returned a very low $p$ value of $1.6 * 10^{-8}$, indicating that a trend exists. The detachment rate, $\beta$, was of the order $\mathcal{O}(10^{-2})$, giving a characteristic timescale between 20 and 77 min for detachment rates for the values displayed in Table S1. It had a weaker, negative, size correlation, with a Pearson's r coefficient of $-0.75$, and a relatively higher Mann-Kendall p-value of $p = 2.1 * 10^{-5}$. However, as neither the Pearson's correlation nor the Mann-Kendall test take errors in the fit parameters into account, we sought to analyze the trend in $\beta$ further.

We extended our analysis of the detachment rate, $\beta$, by examining the standard error in the fit value. For almost all size bins, $\beta > 0$ was found with high statistical significance ($p < 0.05$). The only exception was for groups of radii of $R = 5.45 \pm 0.11$ µm, where $p = 0.15$. We can therefore conclude that a non-zero detachment rate is statistically likely. However, we noticed that the detachment rate for all groups larger than 10 microns in radius was quite flat. We averaged the detachment rate for $R > 10$ microns, finding that in this zone, $\bar{\beta} = 0.019\text{min}^{-1}$. We then used a two-sided t-test to compare the fit values of $\beta$ for each size class with this average value. We found that this test failed to reject the null hypothesis at a $\alpha = 0.05$ significance level for 19/22 size classes: $R = \{5.1, 9.5, 15.6\}$ microns.

To understand how a constant $\beta$ value might affect the timetrack fit, we performed fits to our model where we restricted the values of $\beta$ and $\Gamma$. First, we held $\beta$ to be a constant equal to the net average value $\bar{\beta} = 0.025\text{min}^{-1}$ from the fits to the timetracks above, so that the only free parameter was the collision and attachment kernel, $\Gamma$. We show the result of these fits in Figure S2b. When compared to fits where $\beta$ was allowed to vary, we found that the sum of the root mean square of the residuals was consistently higher for all size classes (Figure S2d). However, on average, the sum of the rms residuals was about 130% of the value of the residuals from the full fit, indicating that the size-dependence of the detachment rate is only mildly important. By comparison, allowing $\beta$ to vary, but fixing $\Gamma$ to be the average of the fits from above, the sum of the root-mean-square of the residuals is on average about 270% higher.

In summary, we did not find enough statistical evidence to conclude that the particle detachment rate, $\beta$, is size-dependent. For the remaining regressions and analysis, we therefore proceeded using a constant $\beta$ value.

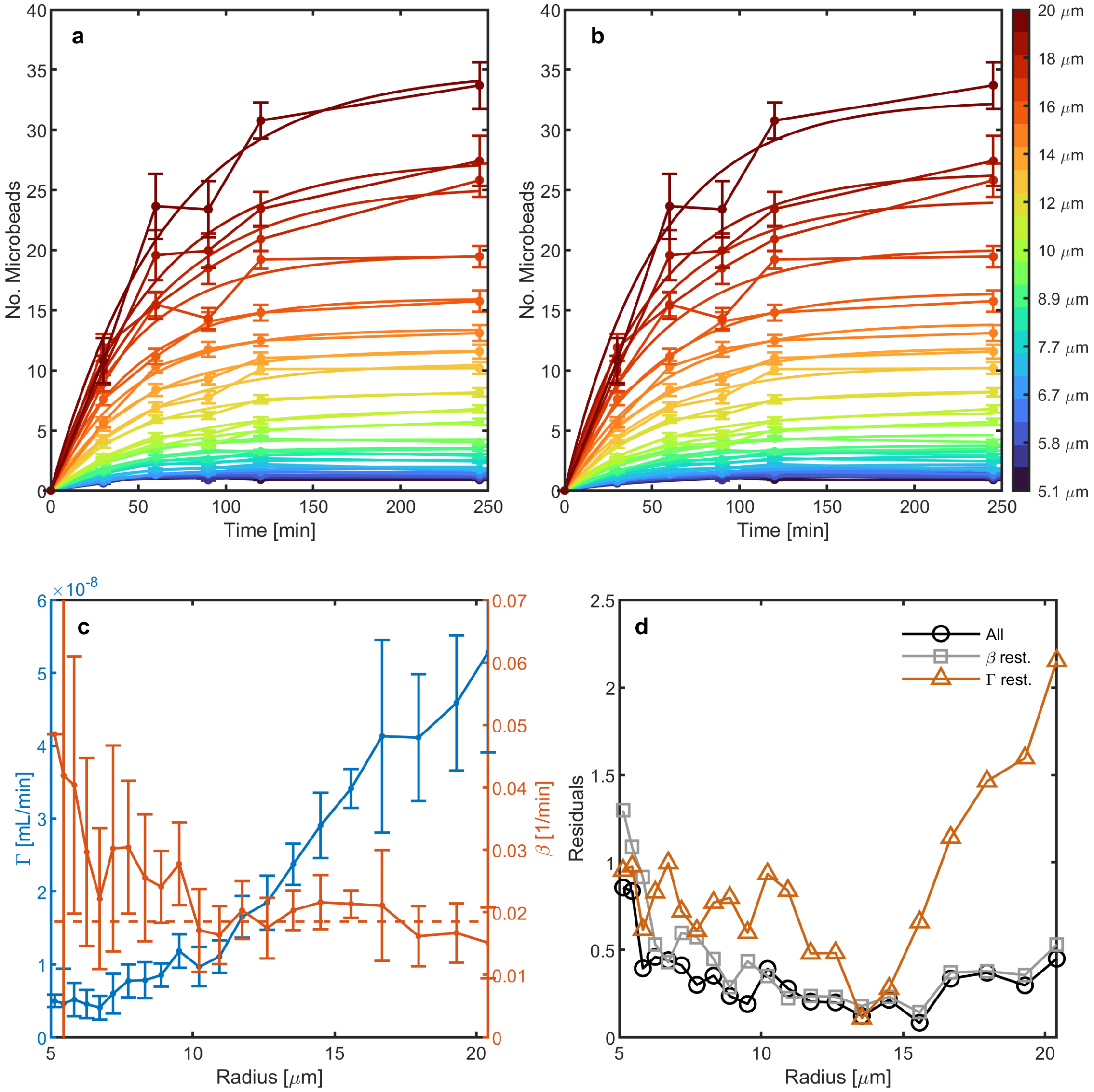


**Figure S2: Average number of microbeads tracked over time for multicellular groups of various size classes.** (a) Colors correspond to size bins. The line fits are regression from Equation 12, allowing both $\beta$ and $\Gamma$ to vary. Errorbars are one standard deviation. (b) Fits to the same data where $\beta$ is held constant. (c) Fit parameters from (a) showing the size dependence of both $\Gamma$ (blue, left axis) and $\beta$ (red, right axis). The dashed horizontal line is the value $\bar{\beta} = 0.019\text{min}^{-1}$, used for fits in (b). Errorbars represent 2 standard errors in the fit value. (d) Sum of the root mean square of the residuals for each size class timetrack.

| $R$ | $\beta$ | $\Gamma$ |
|---|---|---|
| 5.0 | 0.050 | $5.16 * 10^{-9}$ |
| 9.3 | 0.025 | $1.01 * 10^{-8}$ |
| 15.7 | 0.020 | $3.49 * 10^{-8}$ |
| 22.1 | 0.013 | $5.29 * 10^{-8}$ |

**Table S1:** Fitting parameters for timetracks of different multicellular group sizes. R, size class measured as radius in μm; $\beta$, particle detachment rate measured as $\text{min}^{-1}$; $\Gamma$, measured as $\text{cm}^3/\text{min}$.

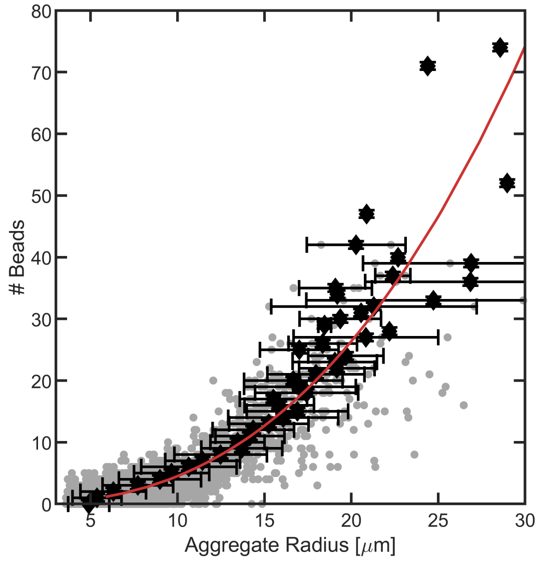


**Figure S3: Number of microbeads attached per multicellular group as a function of multicellular group size** Gray points are individual instances. Black diamonds represent avaverage number of beads for a size bin of a group radius. Horizontal error bars represent standard deviation for radii in the bin, vertical error bars are an estimated error from hand counting. Red line is a power law regression via equation 25. Data are replotted from Figure 2b and represent $N = 3150$ individual data points from one biological replicate.

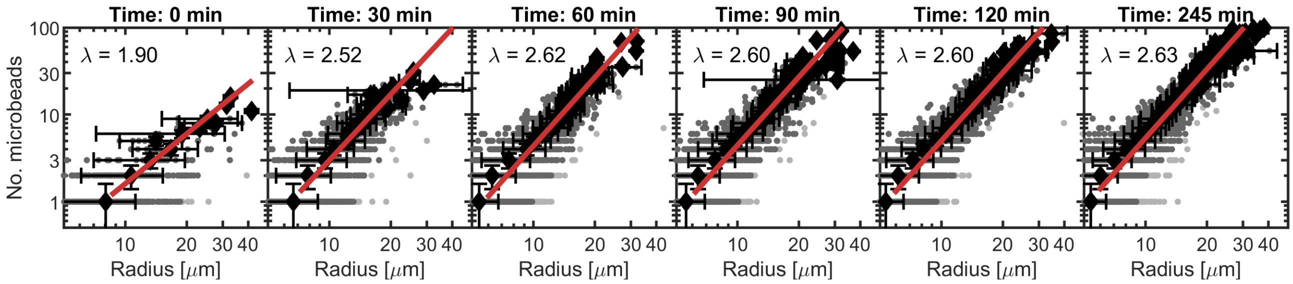


**Figure S4: Number of microbeads attached per multicellular group as a function of multicellular group size sampled over time**. Black diamonds represent averages in each bin. Horizontal error bars are the standard deviation for radii in the bin, vertical error bars are an estimated error in counting particles. Red line is a power law regression. The number of datapoints reported in each panel are: $N = \{1893, 2239, 3442, 3152, 3939, 6167\}$

#### Encounter rate regressions as a function of size

We next sought to measure the mean-field encounter rate as it varied with size. From Equation 12, given no size dependence for the parameter $\beta$, the only size-dependent parameter is the attachment-modified encounter kernel $\alpha\Gamma$. Because of this, and since encounter rate theory of the turbulence kernel both well above and well below the Kolmogorov microscale are power-law relationships, we used a power-law regression to fit the number of microbeads attached as a function of multicellular group size. In Figure 2, we show this regression on a log-log-scale. Additionally, here we show a linear-scale version of this regression in Figure S3.

### Varying incubation, concentration, and shaking

#### Effect of incubation interval on empirical measurement of encounter kernel

In Figure S4, we show the number of microbeads attached to each multicellular group for each timepoint, as a function of the multicellular group size. We found that after about 60 minutes of incubation, there was relatively little change in the nature of power law fit exponent between timepoints, indicating that a steady-state scaling relationship had been reached.

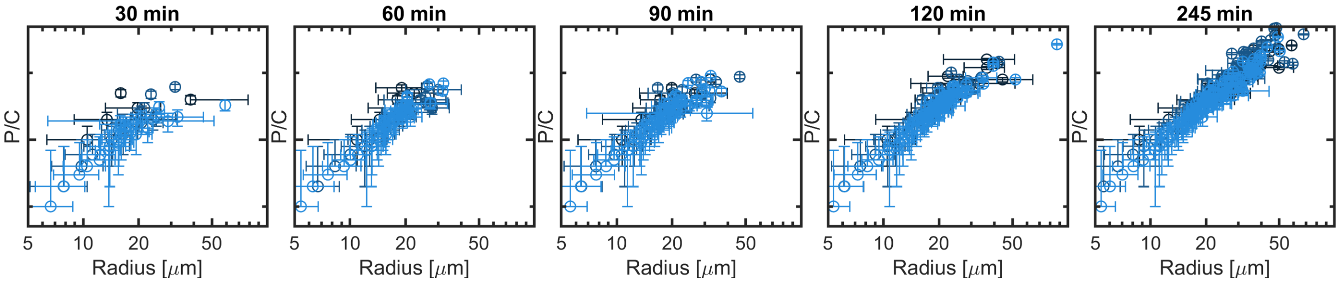


**Figure S5: Number of microbeads attached per multicellular group per unit concentration as a function of group size sampled over time**. $P/C$ is the number of microbeads attached per group, divided by the nominal microbead concentration, for three different concentrations of microbeads. From darkest to lightest blue, nominal concentrations are $C = \{10^6, 5 * 10^6, 10^7\}$ beads per milliliter. For clarity, we show the average group radius for groups with $P$ microbeads attached, with errorbars. Horizontal error bars are the standard deviation for radii in the bin, vertical error bars are an estimated error in counting microbeads. The number of groups shown in each panel for concentrations $\{10^6, 5 * 10^6, 10^7\}$ is as follows: $T = 30$min: $N = \{2042, 2025, 2239\}$, $T = 60$min: $N = \{2286, 3130, 3442\}$, $T = 90$min: $N = \{2663, 2868, 3152\}$, $T = 120$min: $N = \{2837, 2790, 3939\}$, $T = 245$min: $N = \{3422, 5136, 6167\}$.

#### Changing bead concentration

Our attachment/detachment model predicts that the concentration of beads has a linear relationship with the number of beads attached. To test this prediction, we incubated microbial aggregates with three different bead concentrations of $n_b = \{1 * 10^6, 5 * 10^6, 1 * 10^7\}$ beads/mL. We counted the number of beads attached per aggregate in the usual way. Then, we divided the number of beads attached per aggregate by the nominal bead concentration to obtain a measurement of the relative number of attached microbeads, which we label as $P/C$. In Figure S5 we show that the three different concentrations collapse onto the same curve for all timepoints tested, as expected from our attachment/detachment model.

#### Changing shaking properties

We wanted to understand how changing the turbulence of the fluid changes the phenomenological behavior we observed. We approached this in two separate ways. First, we changed the intensity at which we shook the co-culture of multicellular groups and beads, by changing the angular speed of the orbital shaker between 50 and 150 rpm. For each case, we counted the number of beads per multicellular group in the same manner as described above. In Figure S6, we show regressions generated from these three cases. Each were left to shake at their given intensity for 60 min before sampling. We found that increasing the shaking speed decreased the scaling exponent, as reported in the main text.

Second, we changed the shaker and/or beaker that we used to agitate the culture. As a control, we used a single well of a 24-well plate shaken at 100 rpm, as used for all other microbead experiments. Then, we tested an additional two types of incubation. First, we used a larger culturing flask, containing 15 mL of fluid total, and ran it on the same 100 rpm orbital shaker. Next, we changed the shaker to be a side to side platform shaker, which provides a slightly different type of turbulent environment. The side-to-side shaking speed was set to 107 rpm. We found that using different shakers generally led to different scaling exponents for the number of beads vs. aggregate size. This is expected, as the turbulent properties of a fluid are highly sensitive to the boundary conditions [6]. For example, the frictional area of contact between a fluid and its container changes significantly when filling flasks with different volumes and leads to differences in the energy dissipation rate.

### Poisson distribution fits

We fit a Poisson distribution to the distribution of the number of microbeads attached to multicellular groups in various size classes. We did this by binning multicellular groups into logarithmically-spaced size bins. For each size bin, we generated a histogram of the number of multicellular groups with $p$ microbeads attached. We normalized this histogram to produce a probability density function (PDF): each bin was normalized like $v_i = c_i/(N * w_i)$, where $c_i$ is the number of counts in the bin, $w_i = 1$ is the width of the bin, and $N = \sum_i c_i$ is the total number of multicellular groups counted. We then produced a predicted PDF from a Poisson distribution with the same mean, $f(k) = \bar{p}^k e^{-\bar{p}}/k!$, where $\bar{p}$ is the average number of beads attached per multicellular aggregate. We overlay a full panel of these histograms and the Poisson predictions in Figure S8.

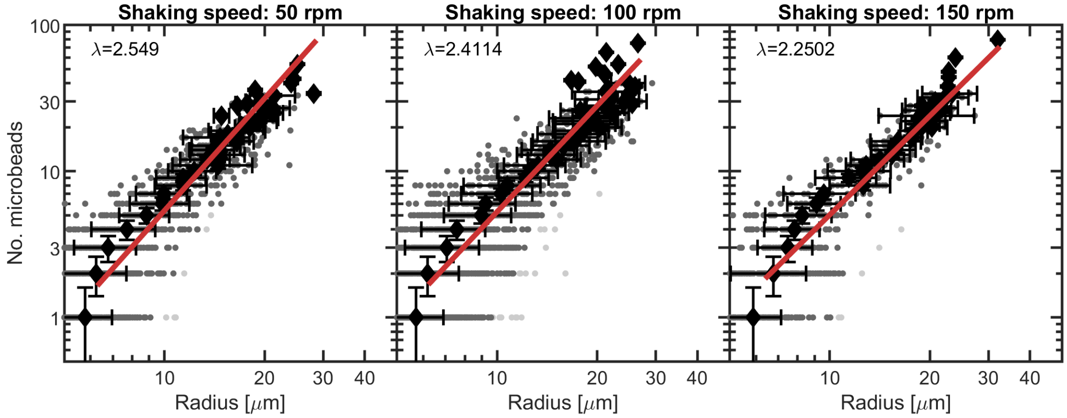


**Figure S6: Distributions of microbeads per bacterial group, measured at different shaking speeds**. All three shaking speeds are with an orbital rotation. Black diamonds represent averages in each bin. Horizontal error bars are the standard deviation for radii in the bin, vertical error bars are an estimated error in counting microbeads. Red line is a power-law regression, and the power $\lambda$ is displayed in the panel. For the three panels, the number of groups counted was $N = \{1472, 2666, 781\}$.

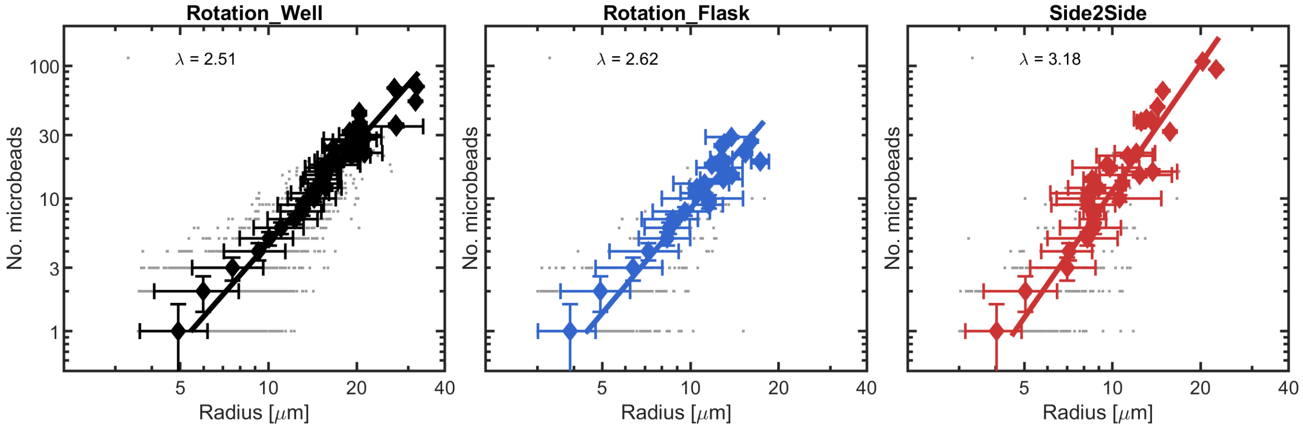


**Figure S7: Distributions of microbeads per bacterial group, measured for different shaking paths (orbital vs. linear) and different culture chambers.** Left: rotation well is an orbital shaker at 100 rpm and a 24-well plate with 0.5 mL of culture. Middle: rotation flask is an orbital shaker at 100 rpm and a 50 mL flask with 15 mL of culture. Right: Side to Side is a linear shaker at 107 rpm and a 24 well plate with 0.5 mL of culture. Diamonds represent averages in each bin. Horizontal error bars are the standard deviation for radii in the bin, vertical error bars are an estimated error in counting microbeads. Overlaid lines are a power-law regression, and the power $\lambda$ is displayed in text in each panel. For the three panels, the number of groups counted was $N = [4189, 1594, 1250]$.

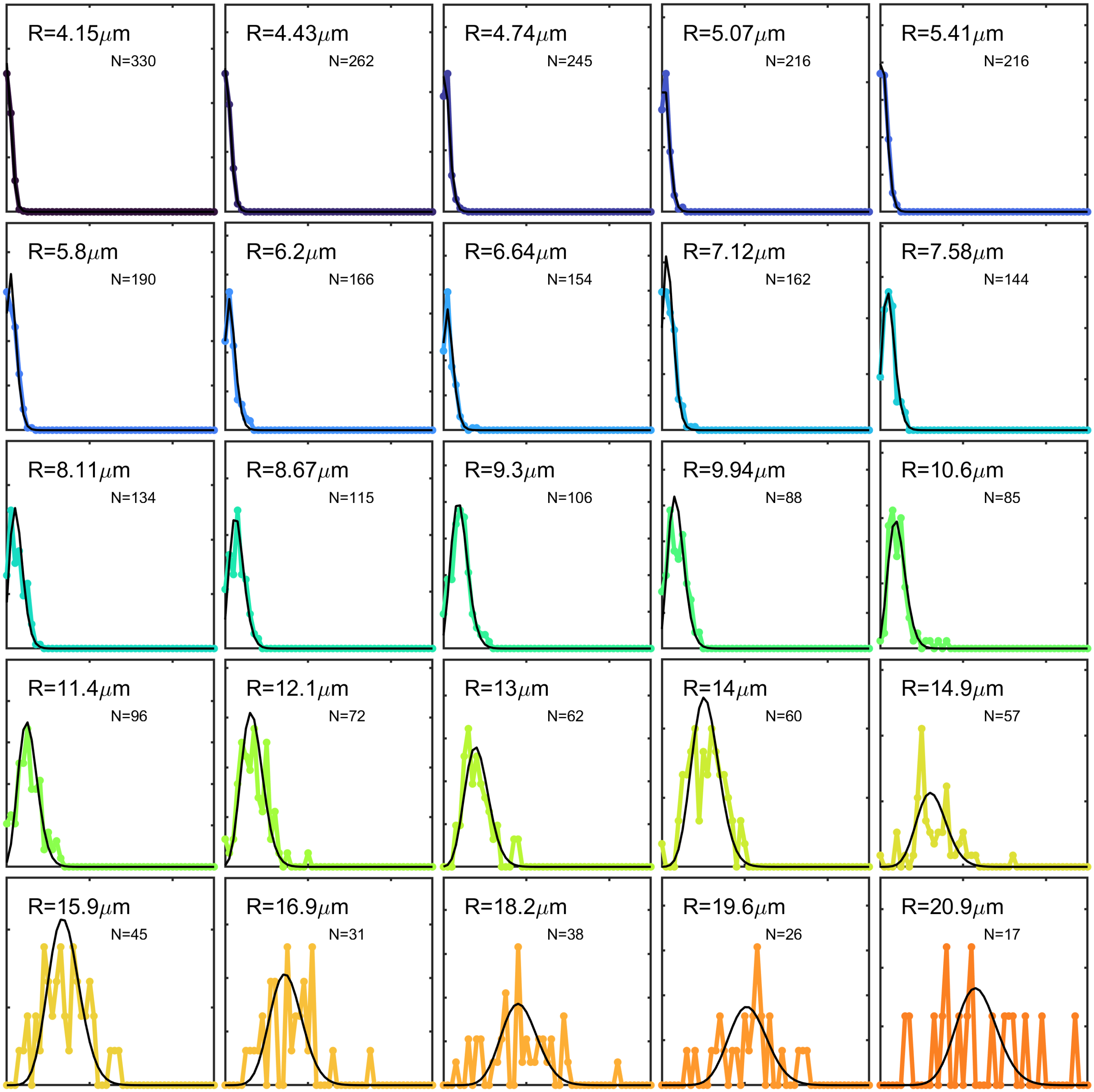


**Figure S8: Poisson distribution matches microbead number histograms for a wide range of multicellular group sizes**. Probability density functions of the number of microbeads attached per multicellular group for groups in different size classes, overlaid with Poisson distribution expectations. The x-axis ranges from 0 to 50 µm in each panel. The y-axis lower limit is always 0, the upper limit is variable to capture the entire distribution. Data are replotted from Figure 2. Text in each panel indicates the average radius of the groups in each bin $R$, and the number of groups in each histogram $N$. We only show bins where $N > 10$.

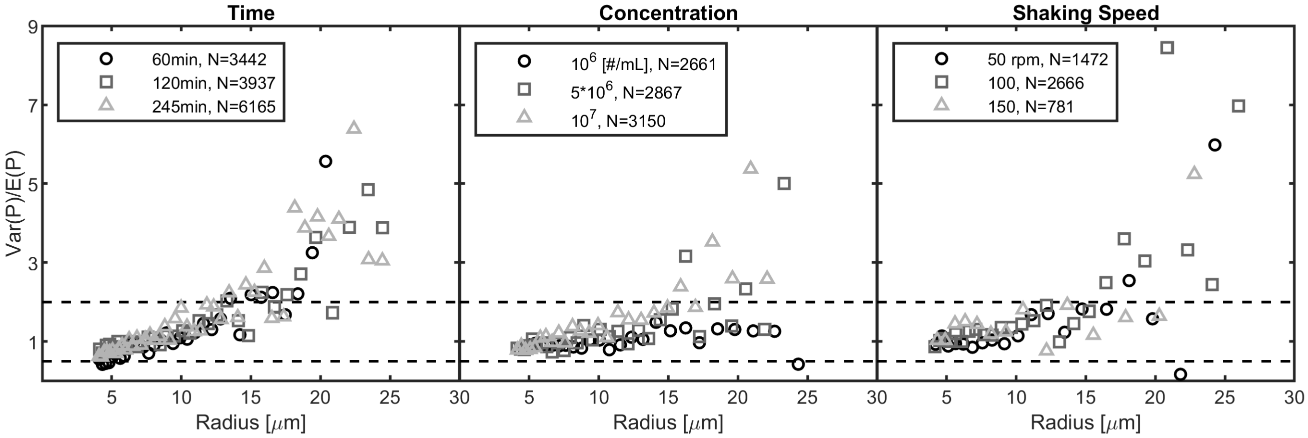


**Figure S9: Level of agreement with Poisson predictions for a wide range of treatments.** Plotted is the variance in the number of microbeads attached per group, divided by the mean number of microbeads attached per group, ($s = \sigma^2/\langle P \rangle$) after binning groups by size. The region between the horizontal dashed lines is where $0.5 < s < 2$. Left: 3 different timepoints, where the concentration of microbeads was $10^7$ beads per mL. Center: 3 different concentrations of bead incubations, all sampled at $T = 90$min. Right: 3 different shaking speeds on an orbital shaker after 60min incubation with $10^7 \text{mL}^{-1}$ bead concentration. Legends indicate the total number of groups counted in each treatment.

#### Measuring agreement with and deviation from Poisson predictions

We measured the degree of agreement with Poisson predictions by comparing the measured mean number of microbeads to the variance in the number of microbeads for binned size classes. We created a test statistic to quantify this comparison,

$$s(r) = \langle (P - \langle P \rangle)^2 \rangle / \langle P \rangle \tag{21}$$

where $\langle . \rangle$ indicates an average over all the multicellular groups in the size bin given by the multicellular group radius $r$, and $P$ is the random variable of the number of attached microbeads to each multicellular group. Since a characteristic of the Poisson distribution is that the mean and variance have equal values, i.e. $s = 1$, we labeled values of the test statistic that were in the range $0.5 < s < 2$ to agree with the Poisson prediction, and values outside this range to disagree. We found that, for a broad range of incubation times, concentrations, and shaking speeds, the test statistic behaved qualitatively the same (Figure S9). Smaller multicellular groups had low values of $s$, which increased as multicellular groups became larger. The exact size at which $s$ generally exceeded our defined range varied slightly between treatments.

#### Testing low statistical power of various size bins

Because the most significant deviations from the Poisson predictions tended to come from size bins where there were relatively few multicellular groups, we wondered if the discrepancy was caused by low statistical counts. To test this, we first ran a random sampling simulation *in silico*, to test how often a Poisson-distributed sample with $m$ members would fail our test statistic criterion, $0.5 < s = \sigma^2/\mu < 2$. We did this by sampling $m$ instances randomly from a Poisson probability density distribution with mean $\lambda = 10$. We then calculated the sample mean, $\bar{x} = \langle x \rangle = 1/m \sum_{i=1}^{m} x_i$, and sample variance, $\sigma^2 = \langle (x - \bar{x})^2 \rangle$, of the set of $m$ instances. We used these sample statistics to calculate our test statistic, $s = \sigma^2/\bar{x}$. We found that samples of size $m = 2$ failed to adhere to the range $0.5, s < 2$ up to 68% of the time. However, when sampling $m = 10$, this only failed our statistic on average about 14% of the time. By comparison, the fail rate of any size bins within our dataset containing between 2 and 10 members was 86%, indicating that random sampling is not the cause of the discrepancy.

#### Connecting multicellular group shape with elevated variance in microbead encounters

We were interested in determining why large multicellular groups generally have a larger variance in the number of microbeads attached to them than the Poisson distribution predicts for their mean number of attachments. We can group reasons for this discrepancy into two general classes: (a) there is a biological cause - for example, it is possible that larger multicellular groups have more variance in their adhesiveness to the microbeads - or (b) there is a physical cause.

One possibility for the physical cause of this variance discrepancy is that large multicellular groups can deviate from spherical shapes. Deviations from sphericity can occur either through growth of the cells within the multicellular group, as cells are oblong and divide via binary fission, or they might occur because multicellular groups can encounter and interact with/adhere to each other. Especially at larger size, multicellular groups are likely to no longer be clonal. As multicellular

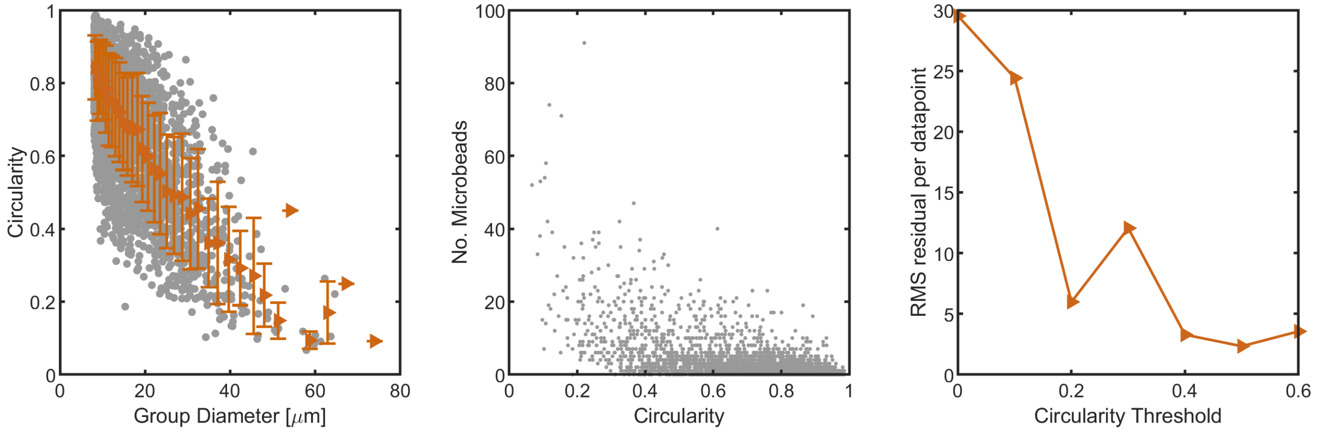


**Figure S10: Loss of circularity in shape is a contributor to elevated variance in microbead attachments.** (a) Circularity decreases with multicellular group size. Gray points are individual measurements ($N = 3150$), orange triangles with errorbars are averages within size bins with standard deviations in circularity. (b) The number of microbeads attached per group vs. the circularity of the group. Gray points are individual group measurements, replotted from S10a. (c) The root-mean-square residual, $r_{rms} = \sqrt{\langle(\bar{p}_i - \sigma_i^2)^2\rangle_i}$, vs. a circularity threshold $C^*$ removing all individuals with circularity less than $C^*$. All data is from the same experiment as Figure 2b.

groups grow in size and concentration, the likelihood that they encounter and attach to one another increases. Such aggregated groups can exhibit strong deviations from sphericity. For example, two spherical multicellular groups, each 10 µm in radius, stuck together such that their centroids are separated by 10 µm, will have a cross-section circularity of 0.69. As multicellular groups adhere into larger aggregates, there become many possible shapes that deviate from sphericity; in other words, for sphericities $< 1$, there are increasingly many possible shapes. As physical encounters are affected by geometry, either a large spread in sphericity, or a simple decrease in sphericity, could contribute to a larger spread in physical encounters. We hypothesized that this physical variance could be contributing to the deviations from the Poisson expectations that we observed for larger group sizes.

To test this hypothesis, we measured the circularity of our groups for one case (we chose $n_b = 1 * 10^7$ beads/mL, $\Delta t = 90$ min, and shaking speed 100 rpm). We found a strong negative correlation between multicellular group size and circularity (Figure S10, supporting the idea that there is more variance in physical shape at larger sizes. We also found that the spread in circularity increased as aggregate diameter increased. Next, we found that the most aspherical groups exhibited the most variance in the number of beads attached. We then applied a range of circularity thresholds to the dataset, whereby we filtered out any groups with cross-sectional circularity below $c$. For each circularity threshold, we measured the size-dependent mean number of attached microbeads $\bar{p}$ and the size-dependent variance $\sigma^2$ and calculated the root-mean-square residual between the mean and variance in each size class ($r_{rms} = \sqrt{\langle(\bar{p}_i - \sigma_i^2)^2\rangle}$). We found that the mean and variance became more similar with more severe circularity thresholds, supporting our hypothesis that deviations from circularity play a contributing role in the elevated variance as compared to the mean rate of attachments for large multicellular groups.

### Measuring number of cells per group

Cellular packing within individual groups was measured for 10 multicellular groups via confocal microscopy. We used cell stain Syto-9 [Invitrogen] at ∼5 µM concentration to label individual cells within 12B01 multicellular group grown on alginate for 36 hrs. Then, we used a Leica point-scanning confocal microscope (see Methods) to image cells within groups. We used a custom segmentation algorithm to segment individual cells within groups. Example segmentation is displayed in Supplemental Figure S11.

#### A description of the 3D segmentation algorithm

In order to segment and count the number of cells in the 3D confocal images, we defined an algorithm that we describe here. It proceeds in parts: Part 0 - Pre-processing, Part 1 - Binarization, Part 2 - Watershedding, Part 3 - Creating Isosurfaces, and Part 4 - Ellipsoid Fitting.

Part 0 - Pre-processing. In the pre-processing step, our goal was to remove extraneous objects (nearby groups and cells not attached to the group of interest) from the scene. First, we blurred the image with an 11x11 gaussian filter with $\sigma = 5$ pixels, followed by a lenient block threshold to binarize the entire 3D image. This ensured that a single group in an image was blurred enough to be counted as one object. We followed this binarization with a round of cubic mask dilation (3 pixels),

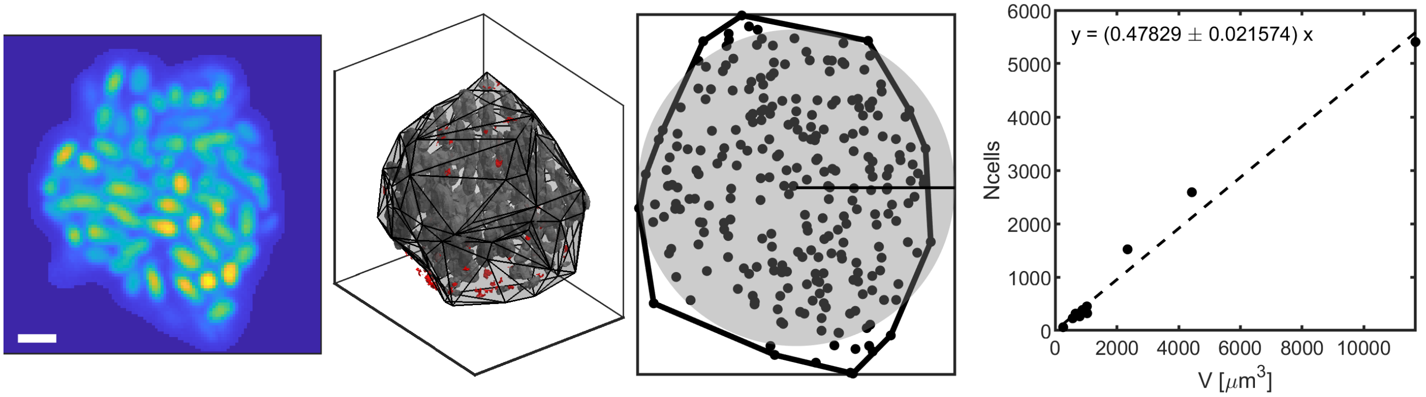


**Figure S11: Cell count is linearly correlated with multicellular group size.** (a) Example confocal z-slice of one bacterial group labeled with Syto-9 fluorescent stain. Scalebar is 2µm. (b) A 3D automated cell segmentation algorithm segmented the cells in each multicellular group. Gray surfaces are segmented cells. In red are shown candidate cells whose ellipsoid fits recorded either negative volumes, negative cell aspect ratios, or aspect ratios above 10. These surfaces were removed from further consideration. (c) Cell centers were projected onto a single z-plane. The 2D convex hull of the points is displayed overtop. A disk of the same cross-sectional area as the convex hull is displayed in gray. The radius of that disk was used to estimate volume. (d) The number of cells is plotted vs. multicellular group volume, showing a linear relationship with $r^2 = 0.98$.

followed by a filling algorithm to fill any extra gaps that had not been caught by the processing so far. We then found all connected components in the mask, and kept only the largest one. We then used this mask to clear any extraneous objects (other cells, smaller groups) from the original image.

Part 1 - Binarization. Next, we sought to binarize individual cells from the background. Since cells become dimmer farther away from the coverslip, we used a local thresholding method known as the Niblack method to binarize each z-slice separately. This method involves computing the average value of a neighborhood surrounding a pixel of interest, and also the standard deviation of the pixels around the pixel of interest. Then, if the pixel of interest exceeds the mean value of the neighborhood by a user-chosen fraction of the standard deviation, the pixel is considered a foreground object. We used a neighborhood of sizes 15x15 pixels, and $\sigma_{\text{thresh}} = 0.5$.

Part 2 - Watershedding. The output from Part 1 is a 3D mask of pixels. However, it is not always able to completely segment neighboring cells from one another. To accomplish this, we used a 3D watershed algorithm that took into account both the local pixel value and its distance to the nearest background pixel to set the watershed lines. We then turned all pixels along the watershed lines into background pixels, effectively splitting the image along these contours. Within each watershed basin, there was thus a single connected component, where all pixels of the object are value 1, and the local background, which has pixels of value 0.

Part 3 - We next determined the surface of each connected component as an isosurface with value 0.5, and listed the vertices of this surface for the next step.

Part 4 - As the bacterial cells are convex objects that can be modeled as ellipsoids, we fit ellipsoids to each connected component vertex list generated from the determined isosurfaces. We later used these ellipsoid fits to measure cellular volume and aspect ratio, and excluded connected components with extreme values for these measurements from the remaining analysis. The parameters we defined as extreme included any negative volume, negative aspect ratio, and any aspect ratio greater than 10. Finally, we saved the information of isosurfaces and ellipsoid fits to file for later use.

Once segmentations were complete, we counted the number of cells in each group and measured the group size. To most accurately emulate how this was measured during brightfield experiments, we first projected all cell centers onto a single z plane, then found the 2D convex hull polygon surrounding these points, then found the radius of a circle with the same cross-sectional area as this convex hull. This radius was then used to obtain an estimated volume of an equivalent sphere. We then used these measurements to generate a regression for the number of cells in a group vs. its volume (Figure S11). We found that the relationship between number of cells and volume was linear: a log-log regression found that the power-law slope was $a = 1.004 \pm 0.04$ (1 sigma). A linear regression returned the fit relationship to be: $N = 0.48V + 24.1$, and had a Pearson's correlation coefficient of $r^2 = 0.993$. We used this regression relationship to estimate the number of cells in all groups henceforth.

### Cell state competition simulations

To simulate competing populations using different strategies to obtain resources from a common pool, we simulated a population of individuals that can grow, divide and die. Inspired by experiments, where multicellular groups are only transient and eventually burst, we assigned individuals to one of two competing cell states. All individuals, no matter their "state", start at the size of a single cell. There is no flux of individuals between the two states. We always seeded competition simulations as occurring between a population of $N = 1000$ cells in state S1, and $N = 1000$ cells in state S2.

**Growth and decay** In both states, individuals grow in size based on encountering food, and reduce in size if they do not encounter enough food. Each individual's biomass changes like $b_i(t + \Delta t) = (\mu_i - \delta)b_i(t)$ where $\mu_i$ represents the growth according to the amount of resources accrued, $\delta$ represents a decay rate, and $i \in [1, N]$ enumerates the individual. All individuals start with a biomass of $b_0$. In State 1, which can be considered the "single-celled state", if an individual's biomass reaches $2.0b_0$, it divides into two individuals of equal biomass (each being $b_0$). In State 2, which can be considered the "multicellular state", individuals never divide, instead continuing to grow in size indefinitely. For either state, if the individual's biomass reaches $0.95b_0$, then it is removed from the population.

**Assigning growth rates** We consider a Monod growth model for growth and division [7]. In this model, growth rate is a saturating function of the amount of food per cell encountered. In other words, the biomass of an individual $i$ increases with rate

$$\mu_i = \frac{\mu_{max}\gamma_i}{\gamma_i + K_m} \tag{22}$$

where $\mu_{max}$ is the maximum possible growth rate, with corresponding doubling time $\tau_m = 1/\mu_{max}$, $K_m$ is the Monod half-saturation constant (which we hold constant between states, and set to be equal to receiving one particle of food per doubling time, $\tau_m$), and $\gamma_i = E_i/N_i$ is the per-capita number of food particles encountered during the minimum doubling time $\tau_m$.

**Fluctuations in food** We ran two separate methods for assigning food, either with or without fluctuations in the amount of food received. Without fluctuations, the amount of food received by each individual is exactly the expected value, being given by $\langle\gamma_i\rangle = \langle E_i\rangle/N_i$ (see below), and can be fractional. With fluctuations, the amount of food received by each individual is randomly assigned from a Poisson distribution with mean and variance given by $\langle E_i\rangle$, and must be an integer value (i.e. the food is discretized into particles).

**Assigning Death Rates** In this model, we held the decay rate $\delta$ constant between the states.

**Calculating the number of resource encounters** The general encounter kernel is

$$\Gamma = \Gamma_D + \Gamma_T + \Gamma_B + ... \tag{23}$$

where the different terms represent the contributions from diffusion, turbulent flow, bouyancy, and more. The number of encounters with food particles during a time period $\Delta t$ that an individual can expect is given by

$$\langle E\rangle = \Gamma c_0 \Delta t \tag{24}$$

where $c_0$ is the concentration of the food particles, and the brackets $\langle ...\rangle$ denote an ensemble average.

Using our empirical measurements of the turbulence kernel, we write the expected number of encounters during a time $\Delta t$:

$$\langle E_i\rangle = Ac_0\Delta t(\theta_i + \theta_0)^\lambda \tag{25}$$

where $A$ is a constant that depends only on properties of the fluid, like the energy dissipation rate and viscosity, and $\theta_0$ is the size of the food particles. The power law constant $\lambda$ was empirically determined to reside between 2 and 3 in our prior experiments; we therefore used this range for $\lambda$ in the remaining simulations. The constant $G_0 = Ac_0$ is the baseline rate of particle encounters; in other words, this is the basal rate of encounters per doubling time of the fastest doubling in the S1 state.

**Running the competition** We initialized a competition experiment with equal proportions of cells in State 1 and State 2. The competition consisted of rounds of selection. Each round, every individual was assigned food according to the food assignment protocol described above. From that, each individual's expected growth rate was calculated. The decay rate was subtracted from this expected growth rate, and the individual's biomass was changed accordingly. Finally, updates in cell division and death were recorded, and the simulation continued.

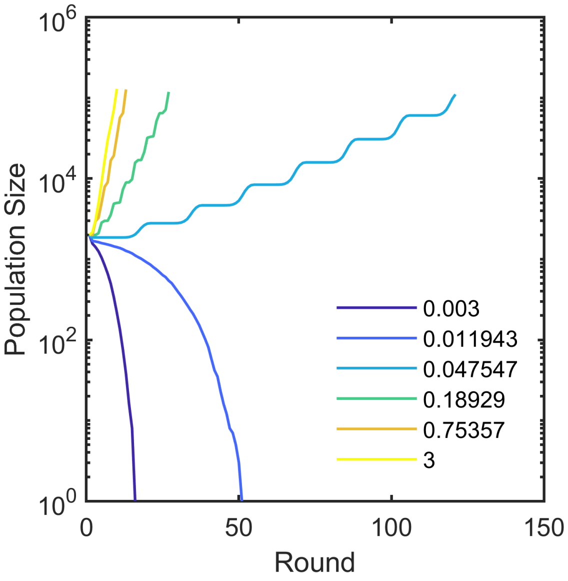


**Figure S12: Population size over simulation time for 6 different simulations, each without any fluctuations in number of resources encountered.** Six different concentrations of resources were chosen according to the average number of particles encountered per maximum cell doubling rate. Concentrations are shown in different colors, from very low concentration (equating to .003 number of encounters per fastest doubling period of the S1 state) to high concentration (3 encounters per basal doubling period of the S1 state).

**Parameter sweeps** We consider the scenario where there is a constant flux of food particles into the volume of interest for the competition. The concentration of food particles sets the baseline rate of encounters between individuals and resources. We chose to sweep resource concentrations such that individuals acquire relatively few resources per their fastest possible doubling time, $\tau_m$; in this regime, cell growth is therefore limited by resource acquisition, especially since the two different strategies S1 and S2 share the same same max doubling rate. We normalized the resource encounters to occur per doubling time $\tau_m$. The explicit range of concentrations we chose swept from between $0.03 \leq G_0 \leq 3$ for a single cell. We chose this range as this was where cells could actually grow. When we pushed the concentration below this level, the population died off (Supplemental Figure S12).

**Including cell swimming** We next sought to include the effect of cell swimming. We model cell swimming as a run-and-tumble pattern without chemotaxis. The nature of this type of swimming is very similar to pure diffusion, with an encounter kernel:

$$\Gamma_S = 4\pi(D_i + D_0)(\theta_i + \theta_0) \tag{26}$$

where $D_0$ is the thermal diffusion constant for a non-swimming particle, and $D_i$ is the "effective" diffusion constant resulting from run-and-tumble motion. A higher $D_i$ means that the cell is swimming more vigorously. We consider the scenario where $D_i \gg D_0$, and so $D_0$ can be considered neglible. Including this swimming term results in the form:

$$\langle E_i \rangle = Ac_0 \Delta t(\theta_i + \theta_0)^\lambda + 4\pi c_0 \Delta t D_i(\theta_i + \theta_0) \tag{27}$$

which we used to assign expected encounters in simulation. We only allowed individuals (of either state) with biomass less than $2b_0$ to swim.

We swept swimming speeds with effective diffusion constants in the range $0.01 \leq D_1 \leq 3000$ μm$^2$/s, thereby sweeping from a range where swimming plays essentially no role in the rate of encounters, to where it plays a dominant role. We show the results from this sweep in Figure 5 of the main text.

To understand what these effective diffusion constants mean for a single swimming organism, we plot the relationship between swimming speed, characteristic run time before re-orientation, and the effective diffusion constant from the random walk in Figure S13. This relationship is well-known [8], as $D = v^2\tau/3$, where $\tau = \sigma/(1-\phi)$, $\sigma$ being the characteristic time between re-orientations, and $\phi$ being the characteristic angle cosine between runs. Here, we use $\phi = 0$, in other words, no correlation between the direction of subsequent runs.

**Growth rate penalties** We next investigated how strong any growth penalties for the multicellular state would have to be in order to recover an advantage for the single celled strategy. We did this by altering the maximum growth rate in the

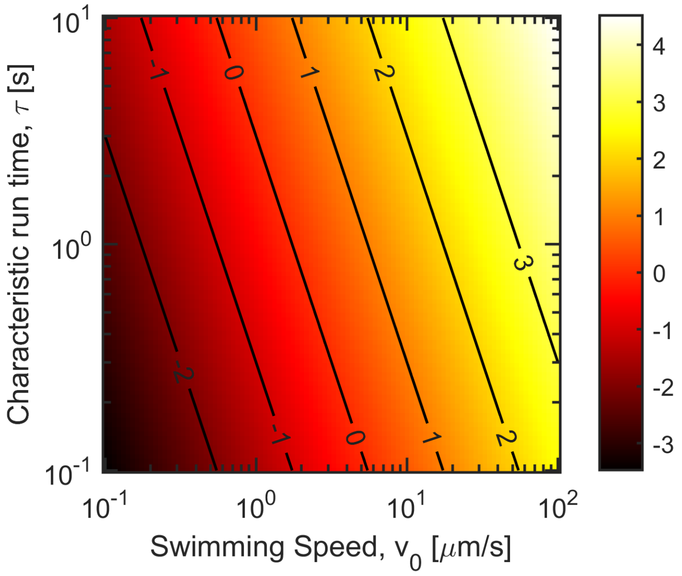


**Figure S13: Effective diffusion constant from swimming.** Color map represents the log of the diffusion constant from $10^{-3}$ to $10^{4}$ μm$^2$/s. Diffusion data are plotted as a function of the swimming speed and the characteristic run time, shown in log10. The labeled contours show swimming strengths of $10^{-2}$ through $10^{3}$ μm$^2$/s, indicating the range swept in simulations.

Monod kinetics, $\mu_{max}$. We assigned a separate $\mu_{max}$ to each of the two cell states (i.e. $\mu_1$ and $\mu_2$), and varied their ratio, such that the fraction $\mu_2/\mu_1$ ranged from 0 (maximum penalty for being in the multicellular state) to 1 (no penalty). We show these results in Figure 5 of the main text.
